# Multi-omics analysis of multiple glucose-sensing receptor systems in yeast

**DOI:** 10.1101/2021.10.07.463466

**Authors:** Shuang Li, Yuanyuan Li, Blake R. Rushing, Sarah E. Harris, Susan L. McRitchie, Daniel Dominguez, Susan J. Sumner, Henrik G. Dohlman

## Abstract

The yeast *Saccharomyces cerevisiae* has long been used to produce alcohol from glucose and other sugars. While much is known about glucose metabolism, relatively little is known about the receptors and signaling pathways that indicate glucose availability. Here we compare the two glucose receptor systems in *S. cerevisiae*. The first is a heterodimer of transporter-like proteins (transceptors), while the second is a seven-transmembrane receptor coupled to a large G protein (Gpa2) and two small G proteins (Ras1 and Ras2). Through comprehensive measurements of glucose-dependent transcription and metabolism, we demonstrate that the two receptor systems have distinct roles in glucose signaling: the G protein-coupled receptor directs carbohydrate and energy metabolism, while the transceptors regulate ancillary processes such as ribosome, amino acids, cofactor and vitamin metabolism. The large G protein transmits the signal from its cognate receptor, while the small G protein Ras2 (but not Ras1) integrates responses from both receptor pathways. Collectively, our analysis reveals the molecular basis for glucose detection and the earliest events of glucose-dependent signal transduction in yeast.

## INTRODUCTION

Most eukaryotic organisms use glucose as the principal source of carbon and energy. Changes in glucose availability result in important metabolic and transcriptional changes that dictate the transition between respiratory and fermentative metabolism [1–4]. Among the best understood systems is that of the yeast *Saccharomyces cerevisiae* (meaning “sugar fungus” and “beer”). Biochemical studies of yeast fermentation led to the discovery of enzymes (meaning “in yeast”) and the founding of biochemistry as a distinct scientific discipline.

While the details of glucose metabolism are well understood, we know comparatively little about changes in signal transduction and cellular metabolism in response to glucose availability. These include changes attributed to glucose binding to cell surface receptors and activation of signaling pathways immediately downstream of the receptor but upstream of glycolysis. In this instance, an increase in glucose is transmitted by two distinct processes (Fig 1). In the first, glucose is detected by a G protein-coupled receptor (GPCR) known as Gpr1 and transmitted through G protein α and β subunits, named Gpa2 and Asc1 respectively [5–13]. This in turn activates the small G proteins Ras1 and Ras2, through the action of guanine nucleotide exchange factors [14–20]. In contrast to *ras2* however, *ras1* has no observed phenotype under standard laboratory growth conditions [21]. Collectively [22–33], these proteins activate the effector enzyme adenylyl cyclase [24, 34–36] leading to an increase in cellular cAMP [9, 26, 37]. This second messenger binds directly to protein kinase A, which goes on to phosphorylate multiple intracellular proteins involved in glucose uptake, metabolism and storage [38–44].

**Fig 1.**
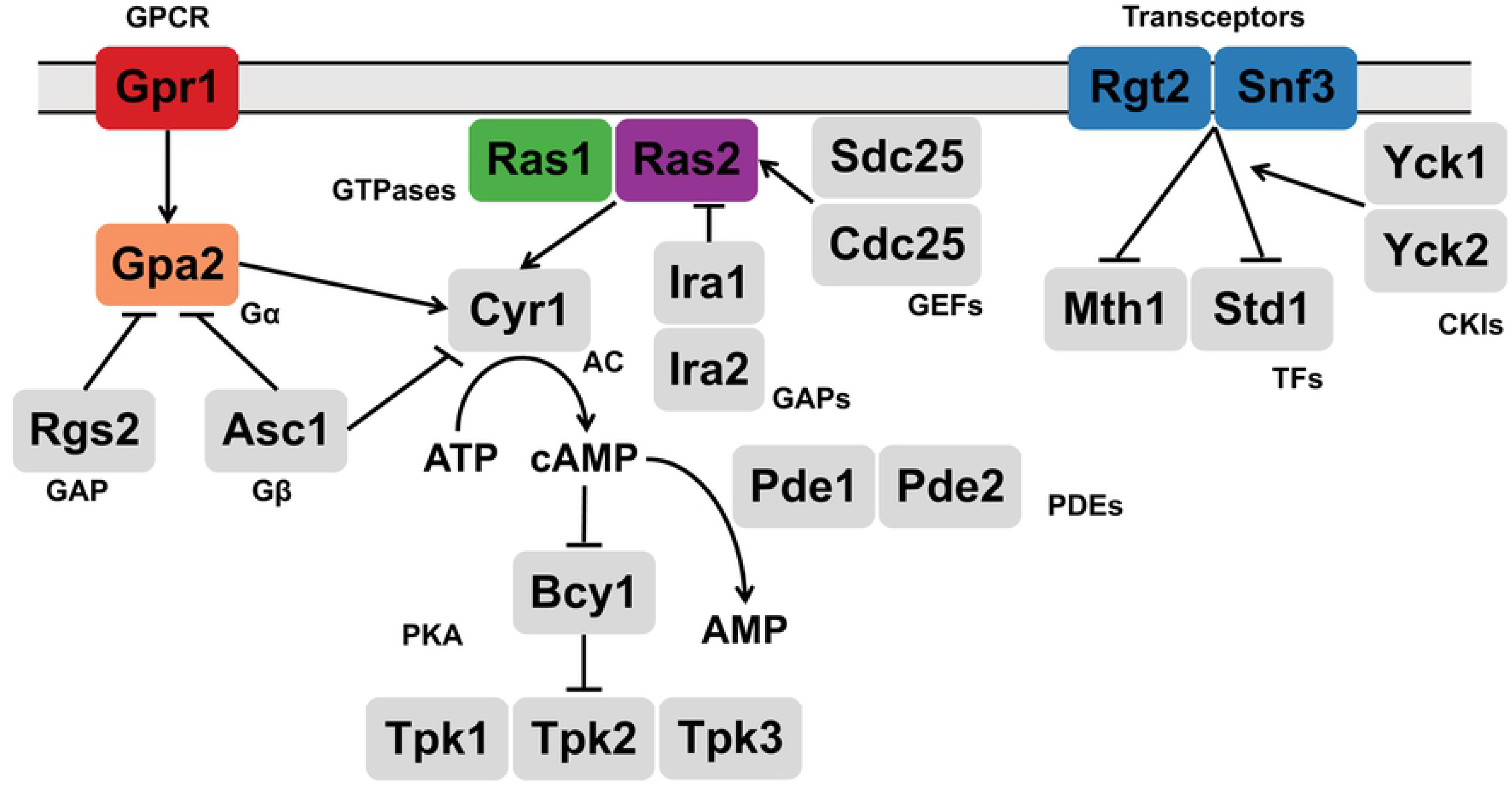
Glucose-sensing pathways in yeast. Two receptor pathways in *S. cerevisiae* respond to glucose availability. Gpr1 transmits its signal through the large G protein Gpa2 [5-7, 9-11], as well as the small G proteins Ras1 and Ras2. The transceptors Snf3 and Rgt2 recruit the protein kinases Yck1 and Yck2 as well as the transcription corepressors Mth1 and Std1 [45–49]. GPCR: G protein coupled receptor; GAP: GTPase activating protein; GEF: guanine nucleotide exchange factor; AC: adenylyl cyclase; PDE: phosphodiesterase; TF: transcription factor; CKI: casein kinase I; PKA: protein kinase A.

The second glucose-sensing system consists of the cell surface proteins Snf3 and Rgt2. Although they resemble glucose transporters, Snf3 and Rgt2 appear to have lost their transporter function and instead serve exclusively as receptor or “transceptor” proteins. Following glucose addition [45–47], Snf3 and Rgt2 recruit the Type I casein kinases Yck1 and Yck2 as well as the transcription corepressors Mth1 and Std1 [48, 49]. Subsequent phosphorylation of these factors results in their ubiquitination and degradation [50–52]; this derepresses genes encoding hexose transporters and promotes the uptake of the newly available sugars [47, 53–62].

Here, we compare the function of the two glucose signaling pathways. In particular, we employ transcriptomics and metabolomics to provide a comprehensive view of the cellular response to glucose. Our analysis reveals new and complementary functions for the two glucose sensing receptors and an unexpected role for Ras2 as an integrator of these two receptor pathways.

## RESULTS

### Unsupervised Principal Component Analysis (PCA)

It is well established that yeast employs two different receptor systems in response to glucose. To investigate the impact of each receptor system, we used untargeted transcriptomics and metabolomics on wildtype cells and mutants lacking the GPCR Gpr1, the large G protein Gpa2, the small G proteins Ras1 or Ras2, or the transceptors Snf3 and Rgt2, under high or low glucose conditions. Log phase wildtype and mutant cells (all prototrophic) were grown for 1 hour in low (L, 0.05%) glucose, then divided and either left untreated or treated with high (H, 2%) glucose for 2 minutes (metabolomics) or 10 minutes (transcriptomics). These time points were selected based on prior data, showing an early and transient spike of cAMP and a subsequent induction of genes within 10 minutes of glucose treatment [1, 13].

Principal Component Analysis (PCA) is an unsupervised multivariate analysis method useful for the visualization of the relationship between observations and variables. When applied to our transcriptomics data, PCA revealed good differentiation based on the proximity of data points for a given treatment and genotype (S1A Fig). This analysis revealed that PC1, which aligns primarily with treatment, accounts for 89% of variance while PC2, which aligns primarily with genotype, represents 4% of variance. Thus, the first 2 components explained 93% of the variance. For metabolomics, the first 2 components explained 50% of the variance (S1B Fig). With the exception of *ras1,* the mutants were distant from wildtype in both measurements. While *gpr1* aligned closely with *gpa2, snf3 rgt2* was on the opposite side of wildtype. The *ras2* mutant was located between the two receptor mutants. These measures indicate distinct effects of the two receptor systems, and a potential role for Ras2 in both.

### Glucose sensing in wildtype cells

Glucose has multiple and complex effects on metabolism and gene expression. To validate our approach, we first performed pathway enrichment analysis, comparing high and low glucose in wildtype cells. For transcriptomics we used the ClusterProfiler package in R [63] and performed gene set enrichment analysis (GSEA) with the Kyoto Encyclopedia of Genes and Genomes (KEGG) database [64–66]. As expected, perturbed pathways were mainly associated with carbohydrate, amino acids, lipids and nucleotide metabolism as well as transcription, ribosome, replication, and cell cycle pathways (See Table 1 in [13], reproduced in S1 Table). These pathways are important for cell growth following the addition of glucose [67]. For metabolomics we used MetaboAnalystR, which integrates the results of Mummichog and GSEA, to produce the combined p-values reported for each pathway (See Table 1 in [13] reproduced in S1 Table) [68–70]. We identified enrichment in six pathways associated with metabolism of carbohydrates, amino acids, and lipids, which is consistent with our transcriptomics analysis. Below we elaborate on how the two receptor systems function individually and in relation to one another.

**Table 1.** Single- and multi-omics integration results for *gpr1*. First block shows GSEA for transcriptomics with adjusted p-value <0.05, arranged in ascending order; second block shows MetaboAnalystR pathway enrichment analysis for metabolomics with combined p-value <0.05, arranged in ascending order; third block shows MetaboAnalystR joint pathway analysis with adjusted p-value <0.05, arranged in ascending order, as detailed in Methods.

| Transcriptomics |  | Metabolomics |  | Integration |  |
| --- | --- | --- | --- | --- | --- |
| enriched pathways | adjusted p-value | enriched pathways | combined p-value | enriched pathways | adjusted p-value |
| Oxidative phosphorylation | 0.0082 | Fructose and mannose metabolism | 0.0021 | Oxidative phosphorylation | 5.24E-19 |
| Starch and sucrose metabolism | 0.0082 | Purine metabolism | 0.0034 | Galactose metabolism | 8.60E-13 |
|  |  | Amino sugar and nucleotide sugar metabolism | 0.0075 | Starch and sucrose metabolism | 6.56E-10 |
|  |  | Galactose metabolism | 0.0075 | ABC transporters | 7.22E-08 |
|  |  | Tyrosine metabolism | 0.0090 | Glycolysis or Gluconeogenesis | 1.75E-07 |
|  |  | Glutathione metabolism | 0.0107 | Arginine biosynthesis | 5.76E-05 |
|  |  | Lysine biosynthesis | 0.0177 | Alanine, aspartate and glutamate metabolism | 0.0001 |
|  |  | Arginine biosynthesis | 0.0229 | Purine metabolism | 0.0001 |
|  |  | Butanoate metabolism | 0.0375 | Citrate cycle (TCA cycle) | 0.0001 |
|  |  |  |  | Fructose and mannose metabolism | 0.0003 |
|  |  |  |  | Amino sugar and nucleotide sugar metabolism | 0.0009 |
|  |  |  |  | Cysteine and methionine metabolism | 0.0025 |
|  |  |  |  | Pentose phosphate pathway | 0.0025 |
|  |  |  |  | Nitrogen metabolism | 0.0128 |
|  |  |  |  | beta-Alanine metabolism | 0.0128 |
|  |  |  |  | Glycine, serine and threonine metabolism | 0.0175 |
|  |  |  |  | Pyruvate metabolism | 0.0264 |
|  |  |  |  | Glutathione metabolism | 0.0266 |

### Comparison of glucose signaling by the GPCR and transceptor systems

#### Transcriptomics analysis

We next determined the transcriptional response to high glucose, comparing wildtype cells with mutants lacking the GPCR (*gpr1),* or the two transceptors (*snf3 rgt2)*. In comparison to wildtype, *gpr1* affected pathways related to oxidative phosphorylation as well as starch and sucrose metabolism, both of which are centered on carbohydrate utilization and energy metabolism (Tables 1 and S2). In comparison to wildtype, *snf3 rgt2* affected pathways related to RNA polymerase, ribosome, autophagy and amino acid metabolism, which are centered on nitrogen utilization and translation (Tables 2 and S3). Thus, under high glucose conditions, the GPCR and transceptor pathways primarily regulate carbohydrate and amino acid metabolism, respectively.

**Table 2.**
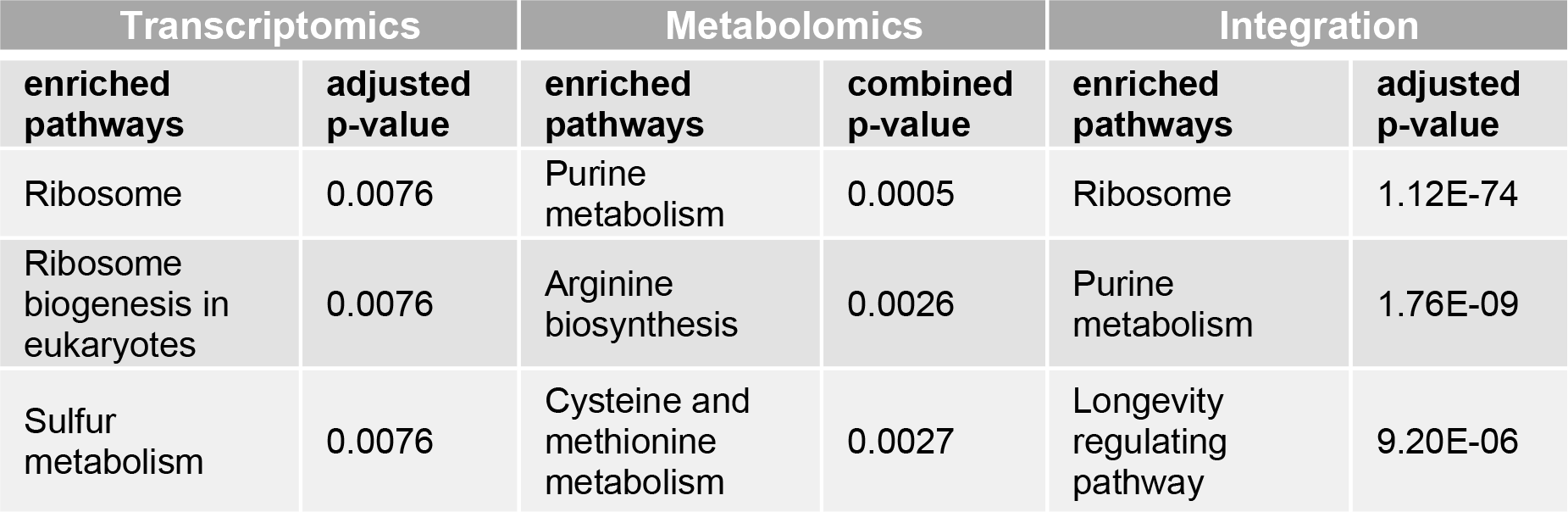
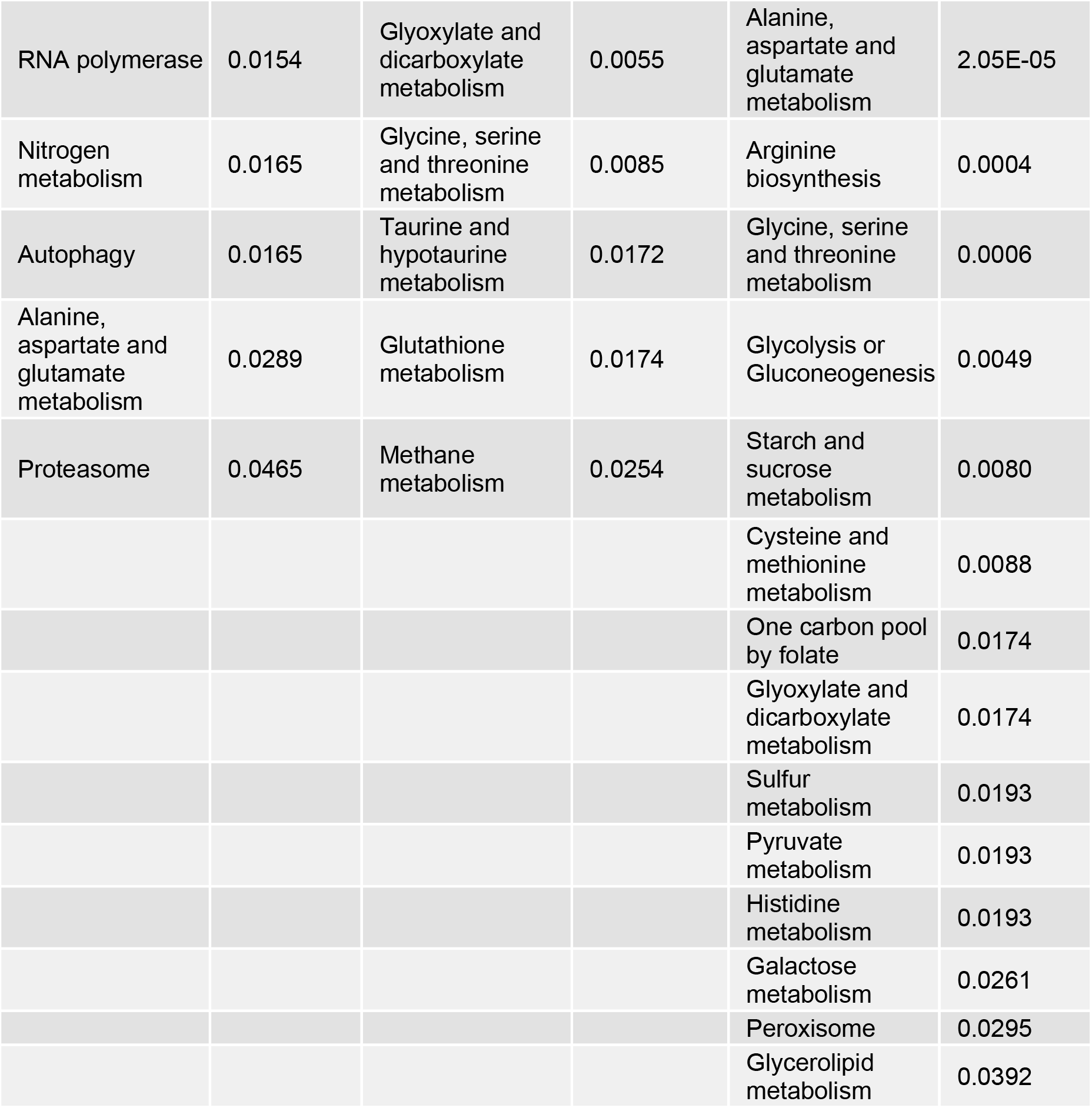
Single- and multi-omics integration results for *snf3 rgt2*. First block shows GSEA for transcriptomics with adjusted p-value <0.05, arranged in ascending order; second block shows MetaboAnalystR pathway enrichment analysis for metabolomics with combined p-value <0.05, arranged in ascending order; third block shows MetaboAnalystR joint pathway analysis with adjusted p-value <0.05, arranged in ascending order, as detailed in Methods.

We then performed over-representation analysis (ORA) for differentially expressed genes (DEGs), comparing *gpr1* vs. wildtype and *snf3 rgt2* vs. wildtype, both under high glucose conditions. In each case we defined the DEGs as having an adjusted p-value <0.05, absolute log2 fold-change value >1 and baseMean >100. Whereas GSEA is a type of functional class scoring that considers a complete list of ranked items (all gene transcripts in this application), ORA considers a thresholded subset of items (DEGs, defined above). In this way, we were able to gain a detailed understanding of how the mutants are similar and how they differ from one another. Fig 2A shows a Venn diagram comparing the specific DEGs for each mutant vs. the wildtype strain (S4 Table). As shown in Fig 2B, DEGs unique to *gpr1* were primarily related to carbohydrate and energy metabolism, consistent with Gpr1’s function as a sensor of glucose availability. DEGs unique to *snf3 rgt2* were mainly related to ribosome, purine, cofactor and vitamin metabolism (Fig 2C). While the two mutant strains had concordant effects on some DEGs (Fig 2D), they had - contrary to our expectations - substantial and opposing effects on a broad set of DEGs primarily related to carbohydrate and amino acid metabolism (Fig 2E). Thus, GSEA and ORA are in agreement, and indicate that the two receptor pathways are largely distinct. When the pathways converge on a shared set of carbohydrate- and amino acid-related transcripts (DEGs), they do so largely in opposition to one another.

**Fig 2.**
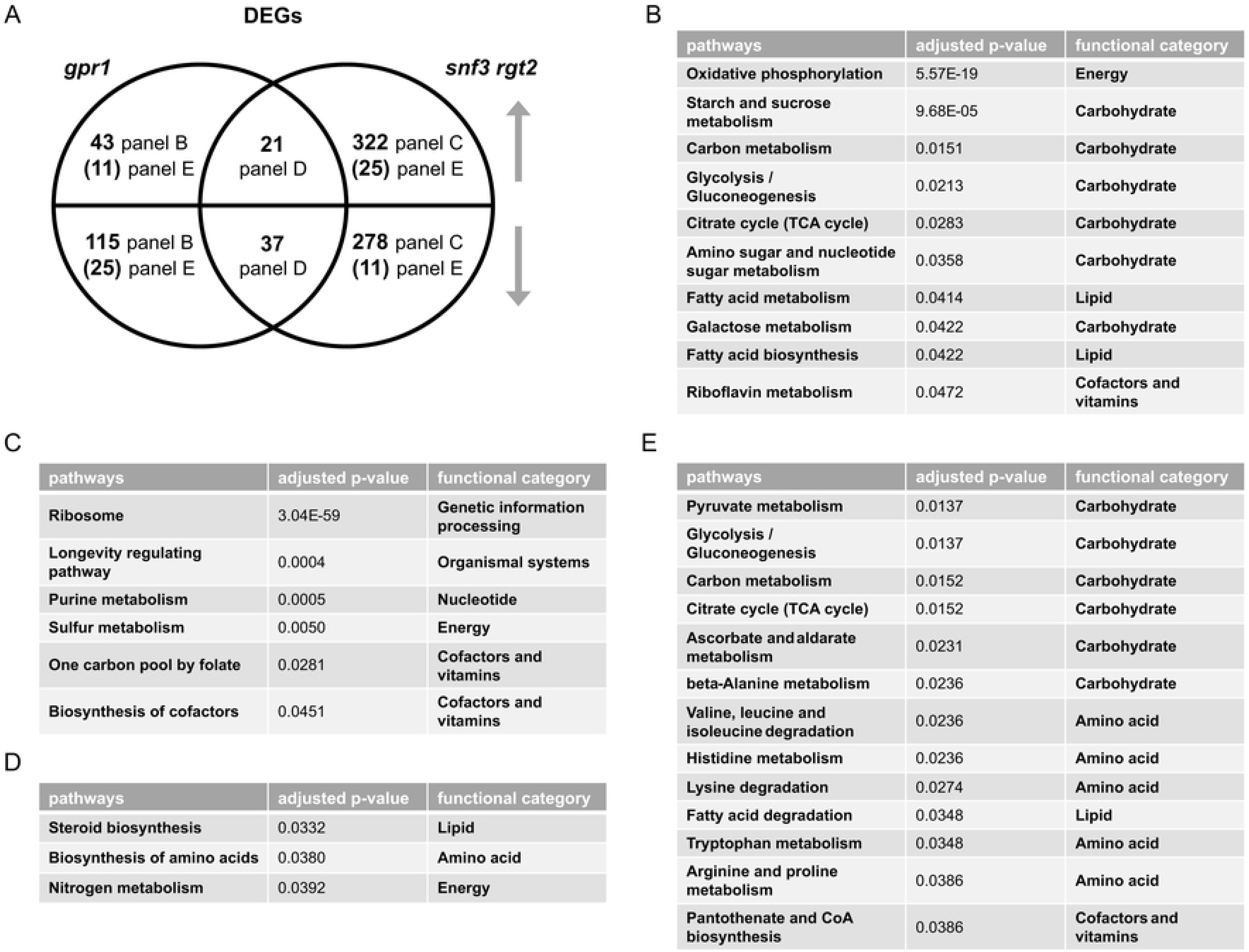
Comparing differentially expressed genes (DEGs) of *gpr1* and *snf3 rgt2*. A) Venn diagram of subsets of DEGs, for *gpr1* vs. wildtype and *snf3 rgt2* vs. wildtype, after glucose addition to 2%. Upper semicircle shows up-regulated DEGs and lower semicircle shows down-regulated DEGs. Numbers in the overlapping region are shared DEGs regulated in the same direction. Numbers in parenthesis are shared DEGs regulated in the opposite direction, and are placed in the area corresponding to the direction of regulation. DEGs used for ORA analysis that are B) unique to *gpr1*; C) unique to *snf3 rgt2*; D) shared and change in the same direction; E) shared and change in the opposite direction. Listed are all pathways and their functional categories with adjusted p-value <0.05.

#### Metabolomics analysis

Gene transcription is regulated by, and in turn regulates, complex metabolic processes in the cell. To better understand the relationship of these two glucose-sensing systems, we examined the role of each receptor type after glucose addition, and did so using untargeted metabolomics. Based on MetaboAnalystR, our mass spectrometry data show that the *gpr1* cells were enriched in nine pathways related to carbohydrate and amino acid metabolism (Tables 1 and S2), while *snf3 rgt2* cells were enriched in eight pathways, including those related to amino acid and purine metabolism, but not central carbohydrate metabolism (Tables 2 and S3). A Venn diagram shows shared and unique metabolites that were significantly perturbed in each strain (Fig 3A and S5 Table). Values were obtained from the output of MetaboAnalystR and represent annotations with adjusted p-value <0.05. These are hereafter referred to as significantly perturbed metabolites (SPMs). ORA revealed that several purine metabolites changed in the same direction (Fig 3B) while a substantial number of carbohydrate metabolites changed in the opposite direction (Fig 3C). As expected, the signals identified and annotated by MetaboAnalyst mirror those obtained using in-house library annotation, developed with data acquired for standards run under the same conditions as the study samples, as well as matching to public databases (PD), as described in our companion manuscript [13], and reported in S6 Table. Subsequent analysis relied on MetaboAnalystR, which is well suited for annotating a large number of signals.

**Fig 3.**
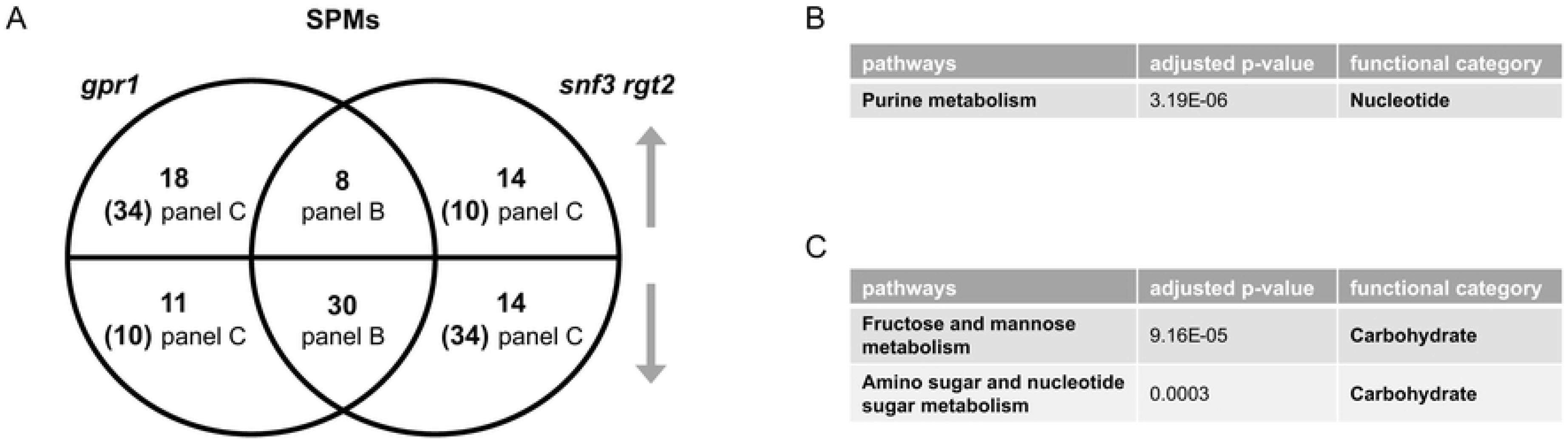
Comparing significantly perturbed metabolites (SPMs) of *gpr1* and *snf3 rgt2*. A) Venn diagram of subsets of SPMs, for *gpr1* vs. wildtype and *snf3 rgt2* vs. wildtype, after glucose addition. Upper semicircle shows up-regulated SPMs and lower semicircle shows down-regulated SPMs. Numbers in the overlapping region are shared SPMs regulated in the same direction. Numbers in parenthesis are shared SPMs regulated in the opposite direction, and are placed in the area corresponding to the direction of regulation. SPMs used for ORA analysis that are B) shared and change in the same direction; C) shared and change in the opposite direction. Listed are all pathways and their functional categories with adjusted p-value <0.05.

#### Integration analysis

Our analysis above shows that the GPCR Gpr1 regulates carbohydrate metabolism while the transceptors Snf3 and Rgt2 regulate ribosome, amino acid, cofactor and vitamin metabolism. Effects that are shared but opposing are primarily related to carbohydrate metabolism; however, these represent only a small subset of the DEGs and SPMs affected by Snf3 and Rgt2. In general, and to our surprise, the glucose transceptors did little to regulate the metabolism of glucose and other sugars. To gain a deeper understanding of the functional relationship between changes in gene transcription and host metabolites, we employed the joint pathway analysis module in MetaboAnalystR, as described previously [13, 69, 70]. In this application, we input all DEGs (transcriptomics) and SPMs (metabolomics) and queried for those over-represented in KEGG. By integrating the data in this manner, we increased the power of our analysis and were able to obtain more information than could be gleaned from transcriptomics or metabolomics alone. Once again, we found that Gpr1 primarily regulates carbohydrate and energy metabolism (Tables 1 and S2) while Snf3 and Rgt2 primarily regulate the ribosome, amino acids, lipids and cofactor metabolism (Tables 2 and S3). Both receptor systems affect genes or metabolites involved in carbohydrates, amino acids and purine metabolites. Thus integration analysis confirms what we observed on the single-omics level: Gpr1 is primarily dedicated to carbohydrate metabolism while Snf3 and Rgt2 work to coordinate other species in response to glucose addition.

To visualize the functional relationship of the two receptor systems, we projected the inputs of our integration analysis onto the pertinent yeast metabolic pathways in KEGG. From this projection it was evident that the two receptor types regulate distinct and complementary processes. Specifically, Gpr1 affects pathways related to carbohydrate metabolism and, within those pathways, a larger number of genes and metabolites compared to Snf3 and Rgt2 (Figs 4A and S2 and Tables 1 and 2). On the other hand, Snf3 and Rgt2 affect pathways related to amino acids and, within those pathways, affect a far greater number of genes and metabolites in comparison to Gpr1 (Figs 4B and S3 and Tables 1 and 2). As presented from the single-omics analysis above, the shared effects on carbohydrate and amino acids were mostly antagonistic while the shared effects on purines were concordant.

**Fig 4.**
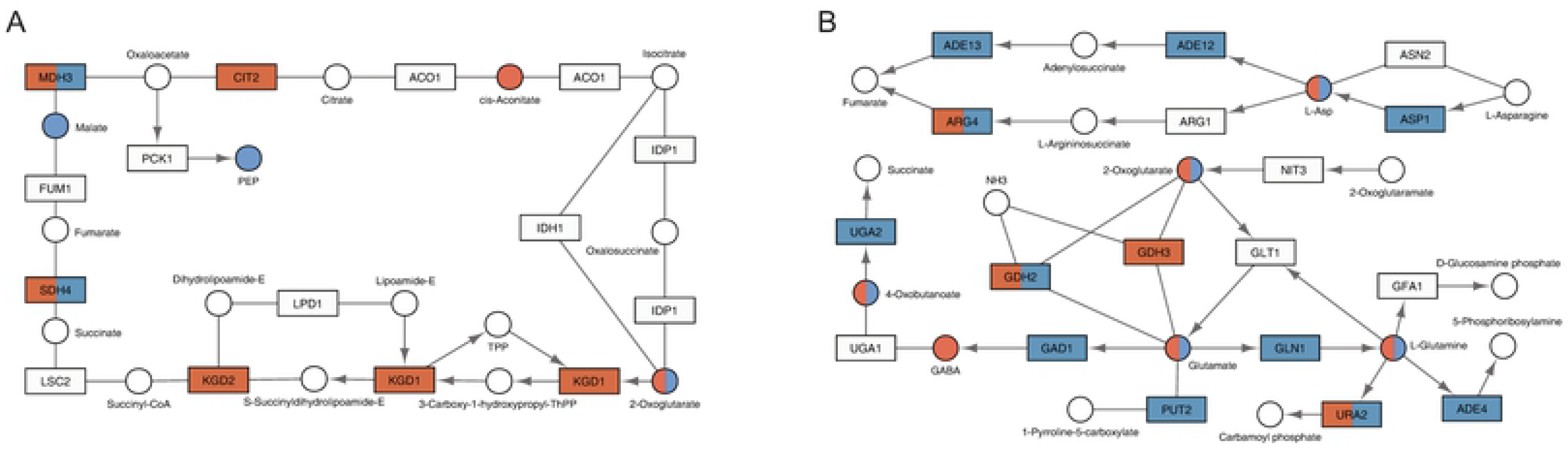
KEGG pathways regulated by *GPR1* or *SNF3* and *RGT2*. Regions of interest in the KEGG pathway are shown with genes displayed as rectangles and metabolites displayed as circles. KEGG compound name for each metabolite is labeled beside the circle. Standard gene names are labeled inside the rectangle. For enzyme complexes, the gene name for the major component is shown followed with an ellipsis. The directions of irreversible enzymatic reactions are shown by the arrows. Reversible reactions are connected by straight lines. DEGs and SPMs are highlighted in red (*gpr1*) and blue (*snf3 rgt2*). Shared DEGs and SPMs are colored half red and half blue. A) as compared with *snf3 rgt2*, *gpr1* affected more components in citrate cycle (TCA cycle, functional category: carbohydrate); B) as compared with *gpr1*, *snf3 rgt2* affected more components in alanine, aspartate and glutamate metabolism (showing aspartate and glutamate specifically, functional category: amino acid).

In summary, our transcriptomics and metabolomics pipeline established a distinct role for each receptor. Two minutes after sugar addition the GPCR and transceptors have opposing effects on many of the same metabolites (Fig 3). After ten minutes however, they confer changes on a largely different set of gene transcripts (Fig 2): whereas the effects of Gpr1 are mostly limited to genes controlling carbohydrate metabolism, Snf3 and Rgt2 affect more diverse species, including genes that are related to amino acids, lipids, ribosome, cofactors and vitamins. Snf3 and Rgt2 do little to alter carbohydrate metabolism, and any changes that do occur are largely in opposition to Gpr1. Such antagonistic effects may allow the cell to fine tune responses and to optimize temporal control of enzyme activities.

### Comparison of glucose signaling through large and small G proteins

Gpr1 acts through a G protein comprised of an α subunit Gpa2 and an atypical Gβ subunit Asc1 [6–9, 12]. Gpa2 in turn activates the small G proteins Ras1 and Ras2, through the action of guanine nucleotide exchange factors [14–20]. Our recent analysis of Gpa2 and Asc1 revealed that they have mostly opposing effects on transcripts and metabolites. When the effects are congruent however, they mirror those observed for their shared activator Gpr1 [13]. To better understand how the receptors transmit their signals in response to glucose addition, we next compared the function of the large and small G proteins (Gpa2, Ras1 and Ras2) using the same analytical pipeline as described above.

#### Transcriptomics Analysis

We began by determining the transcriptional profiles of individual gene deletion mutants by GSEA, as described above. The *ras1* mutant yielded no DEGs, consistent with the lack of phenotype for *ras1* in standard laboratory growth conditions [21]. As shown in Tables 3 and 4 (also S7 and S8 Tables), the *gpa2* mutant affected pathways related to oxidative phosphorylation and ribosome biogenesis. These pathways were likewise regulated by *ras2*. In addition, *ras2* affected RNA polymerase, carbohydrate metabolism and autophagy (Table 4). Overlap between the large and small G proteins was expected given that both are activated by Gpr1 and both are activators of adenylyl cyclase. However, based on the Venn diagram, these mutants had correspondent effects on only a small number of genes and opposing effects on even fewer (Fig 5A and S9 Table). Based on ORA, for the small number of shared DEGs, *gpa2* and *ras2* had mostly concordant effects on processes related to carbohydrate, amino acid and lipid metabolism (Fig 5B). DEGs unique to *ras2* affected a broad spectrum of processes, encompassing all major species in KEGG, including the metabolism of energy, carbohydrates, amino acids, nucleotides, lipids, cofactors and vitamins (Fig 5C). In contrast, DEGs unique to *gpa2* affected a small number of processes, related to carbohydrate, energy and lipid metabolism (Fig 5D). Thus, upon glucose addition, Gpa2 regulates carbohydrates and lipids, while Ras2 affects all major categories of metabolic processes.

**Fig 5.**
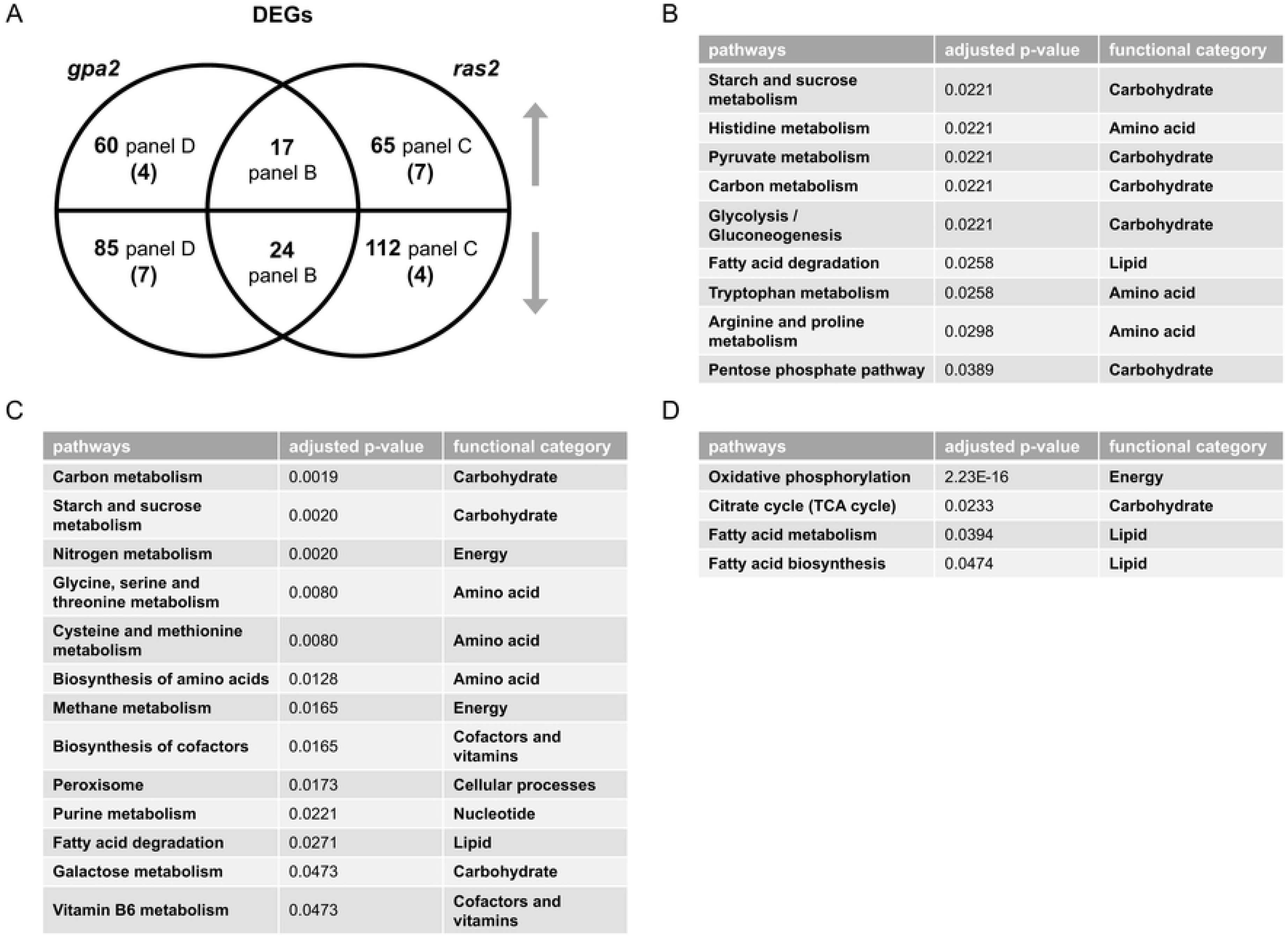
Comparing differentially expressed genes (DEGs) of *gpa2* and *ras2*. A) Venn diagram of subsets of DEGs, for *gpa2* vs. wildtype and *ras2* vs. wildtype, after glucose addition to 2%. Upper semicircle shows up-regulated DEGs and lower semicircle shows down-regulated DEGs. Numbers in the overlapping region are shared DEGs regulated in the same direction. Numbers in parenthesis are shared DEGs regulated in the opposite direction, and are placed in the area corresponding to the direction of regulation. DEGs used for ORA analysis that are B) shared and change in the same direction; C) unique to *ras2*; D) unique to *gpa2.* Listed are all pathways and their functional categories with adjusted p-value <0.05.

**Table 3.** Single- and multi-omics integration results for *gpa2*. First block shows GSEA for transcriptomics with adjusted p-value <0.05, arranged in ascending order; second block shows MetaboAnalystR pathway enrichment analysis for metabolomics with combined p-value <0.05, arranged in ascending order; third block shows MetaboAnalystR joint pathway analysis with adjusted p-value <0.05, arranged in ascending order, as detailed in Methods.

| Transcriptomics |  | Metabolomics |  | Integration |  |
| --- | --- | --- | --- | --- | --- |
| enriched pathways | adjusted p-value | enriched pathways | combined p-value | enriched pathways | adjusted p-value |
| Ribosome biogenesis in eukaryotes | 0.0084 | Purine metabolism | 0.0021 | Oxidative phosphorylation | 3.13E-14 |
| Oxidative phosphorylation | 0.0084 | Fructose and mannose metabolism | 0.0047 | Galactose metabolism | 1.60E-11 |
|  |  | Amino sugar and nucleotide sugar metabolism | 0.0079 | ABC transporters | 1.31E-08 |
|  |  | Galactose metabolism | 0.0079 | Glycolysis or Gluconeogenesis | 8.02E-05 |
|  |  | Glutathione metabolism | 0.0155 | Fructose and mannose metabolism | 8.18E-05 |
|  |  | Tyrosine metabolism | 0.0224 | Starch and sucrose metabolism | 8.18E-05 |
|  |  | Arginine biosynthesis | 0.0234 | Arginine biosynthesis | 0.0012 |
|  |  | Biotin metabolism | 0.0252 | Pentose phosphate pathway | 0.0012 |
|  |  | Aminoacyl-tRNA biosynthesis | 0.0360 | Purine metabolism | 0.0017 |
|  |  | Starch and sucrose metabolism | 0.0396 | Amino sugar and nucleotide sugar metabolism | 0.0073 |
|  |  | Phosphatidylinositol signaling system | 0.0424 | beta-Alanine metabolism | 0.0119 |
|  |  |  |  | Alanine, aspartate and glutamate metabolism | 0.0139 |
|  |  |  |  | Cysteine and methionine metabolism | 0.0149 |
|  |  |  |  | Citrate cycle (TCA cycle) | 0.0221 |

**Table 4.** Single- and multi-omics integration results for *ras2*. First block shows GSEA for transcriptomics with adjusted p-value <0.05, arranged in ascending order; second block shows MetaboAnalystR pathway enrichment analysis for metabolomics with combined p-value <0.05, arranged in ascending order; third block shows MetaboAnalystR joint pathway analysis with adjusted p-value <0.05, arranged in ascending order, as detailed in Methods.

| Transcriptomics |  | Metabolomics |  | Integration |  |
| --- | --- | --- | --- | --- | --- |
| enriched pathways | adjusted p-value | enriched pathways | combined p-value | enriched pathways | adjusted p-value |
| Starch and sucrose metabolism | 0.0065 | Lysine biosynthesis | 1.00E-05 | Galactose metabolism | 1.24E-15 |
| Oxidative phosphorylation | 0.0065 | Glyoxylate and dicarboxylate metabolism | 0.0005 | Starch and sucrose metabolism | 1.68E-11 |
| Ribosome biogenesis in eukaryotes | 0.0065 | Glycine, serine and threonine metabolism | 0.0009 | Glycolysis or Gluconeogenesis | 6.26E-08 |
| Ribosome | 0.0065 | Arginine biosynthesis | 0.0017 | Glycine, serine and threonine metabolism | 4.30E-06 |
| RNA polymerase | 0.0092 | Cysteine and methionine metabolism | 0.0028 | ABC transporters | 1.03E-05 |
| Galactose metabolism | 0.0169 | Taurine and hypotaurine metabolism | 0.0043 | Pentose phosphate pathway | 2.27E-05 |
| Autophagy | 0.0169 | Amino sugar and nucleotide sugar metabolism | 0.0043 | Cysteine and methionine metabolism | 2.27E-05 |
| Meiosis | 0.0332 | Galactose metabolism | 0.0043 | Fructose and mannose metabolism | 0.0002 |
| Spliceosome | 0.0458 | Butanoate metabolism | 0.0095 | Amino sugar and nucleotide sugar metabolism | 0.0002 |
|  |  | Lysine degradation | 0.0098 | Arginine biosynthesis | 0.0003 |
|  |  | Aminoacyl-tRNA biosynthesis | 0.0098 | Purine metabolism | 0.0005 |
|  |  | Starch and sucrose metabolism | 0.0204 | Glyoxylate and dicarboxylate metabolism | 0.0009 |
|  |  | Alanine, aspartate and glutamate metabolism | 0.0372 | Peroxisome | 0.0014 |
|  |  | Fructose and mannose metabolism | 0.0384 | Methane metabolism | 0.0057 |
|  |  | Nitrogen metabolism | 0.0392 | Alanine, aspartate and glutamate metabolism | 0.0057 |
|  |  | Phosphatidylinositol signaling system | 0.0429 | Lysine biosynthesis | 0.0213 |
|  |  |  |  | Nitrogen metabolism | 0.0255 |
|  |  |  |  | Citrate cycle (TCA cycle) | 0.0338 |
|  |  |  |  | Vitamin B6 metabolism | 0.0385 |
|  |  |  |  | Inositol phosphate metabolism | 0.0390 |
|  |  |  |  | Monobactam biosynthesis | 0.0392 |
|  |  |  |  | Tryptophan metabolism | 0.0464 |

#### Metabolomics Analysis

To better understand the relationship of large and small G proteins, we next conducted untargeted metabolomics on the corresponding mutants after glucose addition. Again, we used MetaboAnalystR for pathway enrichment analysis. In agreement with our transcriptomics data, we found that *gpa2* and *ras2* affected several common pathways related to carbohydrate and amino acid metabolism; however, *ras2* impacted a wider variety of amino acid species (Tables 3, 4, S7 and S8). Based on the Venn diagram and ORA analysis, *gpa2* and *ras2* had a large number of shared SPMs that changed in the same direction, most of which were related to carbohydrate metabolism (Fig 6 and S10 Table). In summary, two minutes after sugar addition the *gpa2* and *ras2* strains exhibited a similar metabolic profile (Fig 6). However, after ten minutes, *gpa2* and *ras2* exhibited a different transcriptional profile (Fig 5): while *gpa2* mainly affected transcripts related to carbohydrates and lipids, *ras2* impacted transcripts related to all major categories of metabolic processes. By any measure, the *ras1* mutant yielded no significant differences, at least under the experimental conditions used in this analysis.

**Fig 6.**
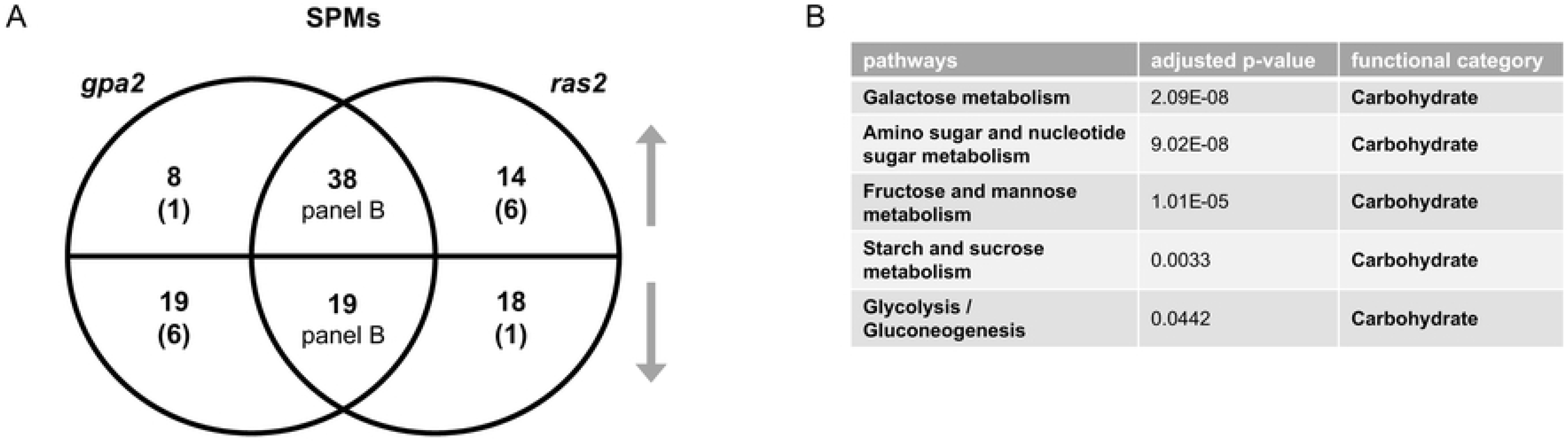
Comparing significantly perturbed metabolites (SPMs) of *gpa2* and *ras2*. A) Venn diagram of subsets of SPMs, for *gpa2* and *ras2* vs. wildtype, after glucose addition. Upper semicircle shows up-regulated SPMs and lower semicircle shows down-regulated SPMs. Numbers in the overlapping region are shared SPMs regulated in the same direction. Numbers in parenthesis are shared SPMs regulated in the opposite direction, and are placed in the area corresponding to the direction of regulation. SPMs used for ORA analysis that are B) shared and change in the same direction. Listed are all pathways and their functional categories with adjusted p-value <0.05.

#### Integration analysis

We then conducted integration analysis using the joint pathway analysis module in MetaboAnalystR. By this method we found that Ras2, like Gpa2, regulated pathways related to carbohydrate, purine and certain amino acids metabolism (Tables 3, 4, S7 and S8). The extent of overlap was particularly evident through integration of the metabolomics and transcriptomics analysis. The *ras2* strain was unique in regulating additional amino acids, as well as lipid and vitamin metabolism (Tables 4 and S8). The *gpa2* strain was unique in regulating oxidative phosphorylation and β-alanine metabolism (Tables 3 and S7). The results obtained using MetaboAnalystR were reflected in the high confidence annotations obtained using our in-house library.

To visualize the functional relationship of Ras2 and Gpa2, we projected the inputs of our integration analysis onto the pertinent yeast metabolic pathways in KEGG. From this visualization, it is evident that the effects of Gpa2 are centered on carbohydrate and energy metabolism, which is shared by Ras2 (Figs 7A and S4). In addition, Ras2 also affects a substantial number of different metabolic species (Figs 7B and S5).

**Fig 7.**
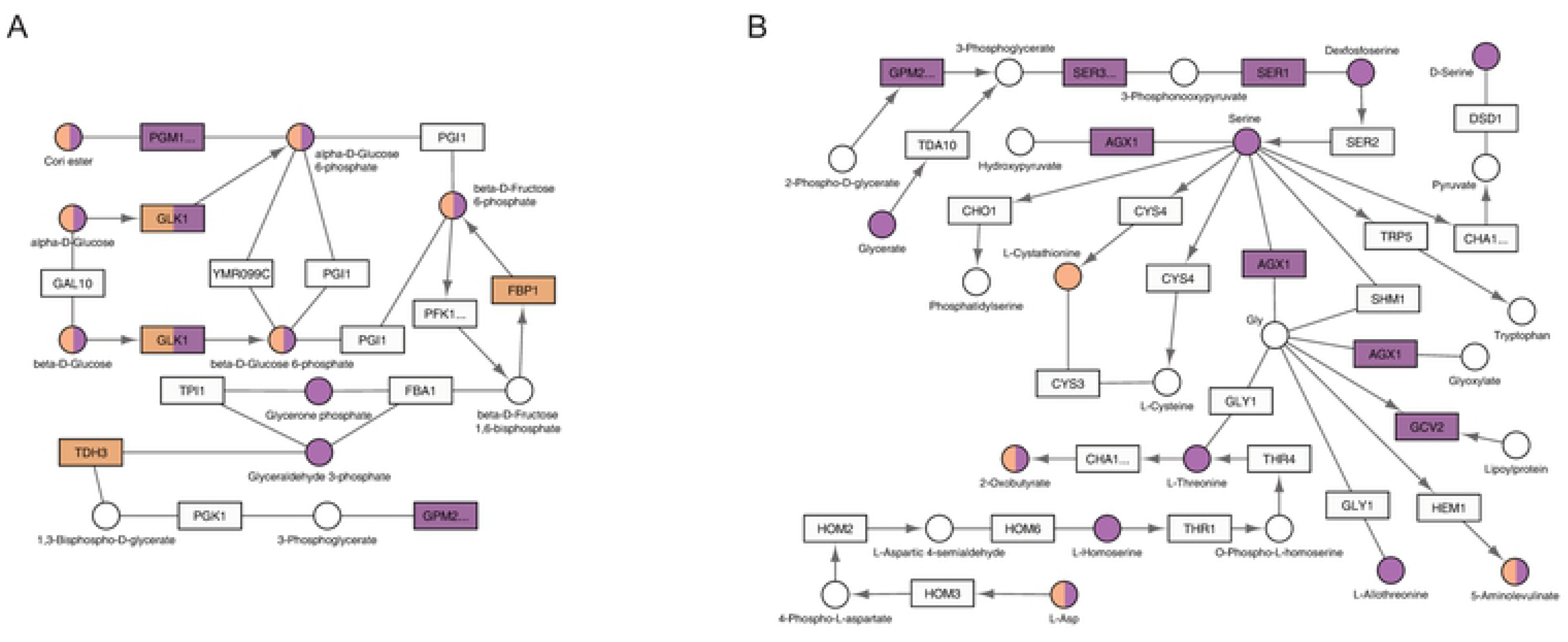
KEGG pathways regulated by *GPA2* or *RAS2*. The relevant part of a specific KEGG pathway is shown with genes displayed as rectangles and metabolites displayed as circles. KEGG compound name for each metabolite is labeled beside the circle. Standard gene names are labeled inside the rectangle. For enzyme complexes, the gene name for the major component is shown followed with an ellipsis. The directions of irreversible enzymatic reactions are shown by the arrows. Reversible reactions are connected by straight lines. DEGs and SPMs are highlighted in orange (*gpa2*) and purple (*ras2*). Shared DEGs and SPMs are colored half orange and half purple. A) both *gpa2* and *ras2* affected components in glycolysis / gluconeogenesis (functional category: carbohydrate); B) as compared with *gpa2*, *ras2* affected more components in glycine, serine and threonine metabolism (functional category: amino acid).

To summarize, we observed three major differences when comparing *gpr1* vs. *snf3 rgt2* and *gpa2* vs. *ras2*. First, the large and small G proteins (Gpa2 and Ras2) had concordant effects on genes and metabolites while the effects of the GPCR (Gpr1) and the transceptors (Snf3 and Rgt2) were largely opposing. Second, Ras2 had considerable effects on carbohydrate metabolism while Snf3 and Rgt2 had little effect on these processes. Third, Ras2, like Snf3 and Rgt2, affected non-carbohydrate-related pathways. However, Ras2 affected far fewer genes and metabolites, as compared to Snf3 and Rgt2. These findings highlight the functional interrelationship of the two receptor systems as well as that of the large and small G proteins.

### Ras2 integrates signals from Gpr1 and Snf3/Rgt2

Above we show that Gpr1 is dedicated to carbohydrate metabolism while Snf3 and Rgt2 primarily control the metabolism of non-carbohydrate species. The downstream G proteins, Gpa2 and Ras2, have concordant effects on carbohydrate metabolism. However, Ras2 affects additional major species that are also affected by Snf3 and Rgt2. Based on these results, we postulated that Snf3 and Rgt2 signal through Ras2. Just as Ras2 acts in synchrony with Gpr1 and Gpa2 to regulate carbohydrate metabolism, we considered if Ras2 also works together with Snf3 and Rgt2 to regulate non-carbohydrate species. Upon examination of the integration analysis presented above, we determined that processes regulated by both Ras2 and Gpa2 are primarily related to carbohydrates (7 pathways shared), and to a lesser extent amino acids (3 pathways shared), as well as purine metabolism (Tables 3, 4, S7 and S8). In comparison, processes regulated by both Ras2 and Snf3/Rgt2 are related to amino acids (4 pathways shared), carbohydrates (4 pathways shared), peroxisome and purines (Tables 2, 4, S3 and S8).

We then quantified DEGs and SPMs regulated by Ras2 as well as by Snf3 and Rgt2. These data are presented as Venn diagrams in Figs 8A and 9A (S11 and S12 Tables). In accordance with our hypothesis, ORA revealed that Ras2 and the transceptors had concordant effects on DEGs related to amino acids, energy, cofactors and vitamins (Fig 8B). While *snf3 rgt2* uniquely affected some DEGs related to purines (Fig 8C), they shared with *ras2* the ability to regulate SPMs related to the same process (Fig 9B). Furthermore, the effect of *ras2* on carbohydrates is not shared by *snf3 rgt2* (Fig 8D). We then performed qPCR to quantify the expression level of genes in wildtype, *ras2* and *snf3 rgt2* before and 10 min after glucose addition. As shown in Fig 10, *ras2* and *snf3 rgt2* have concordant effects on *INO1*, which encodes an Inositol-3-phosphate synthase, and *AGX1*, the product of which catalyzes the synthesis of glycine from glyoxylate; neither enzyme is directly related to carbohydrate metabolism. Thus, multiple lines of evidence indicate that Ras2 and the transceptors share the ability to regulate non-carbohydrate metabolism.

**Fig 8.**
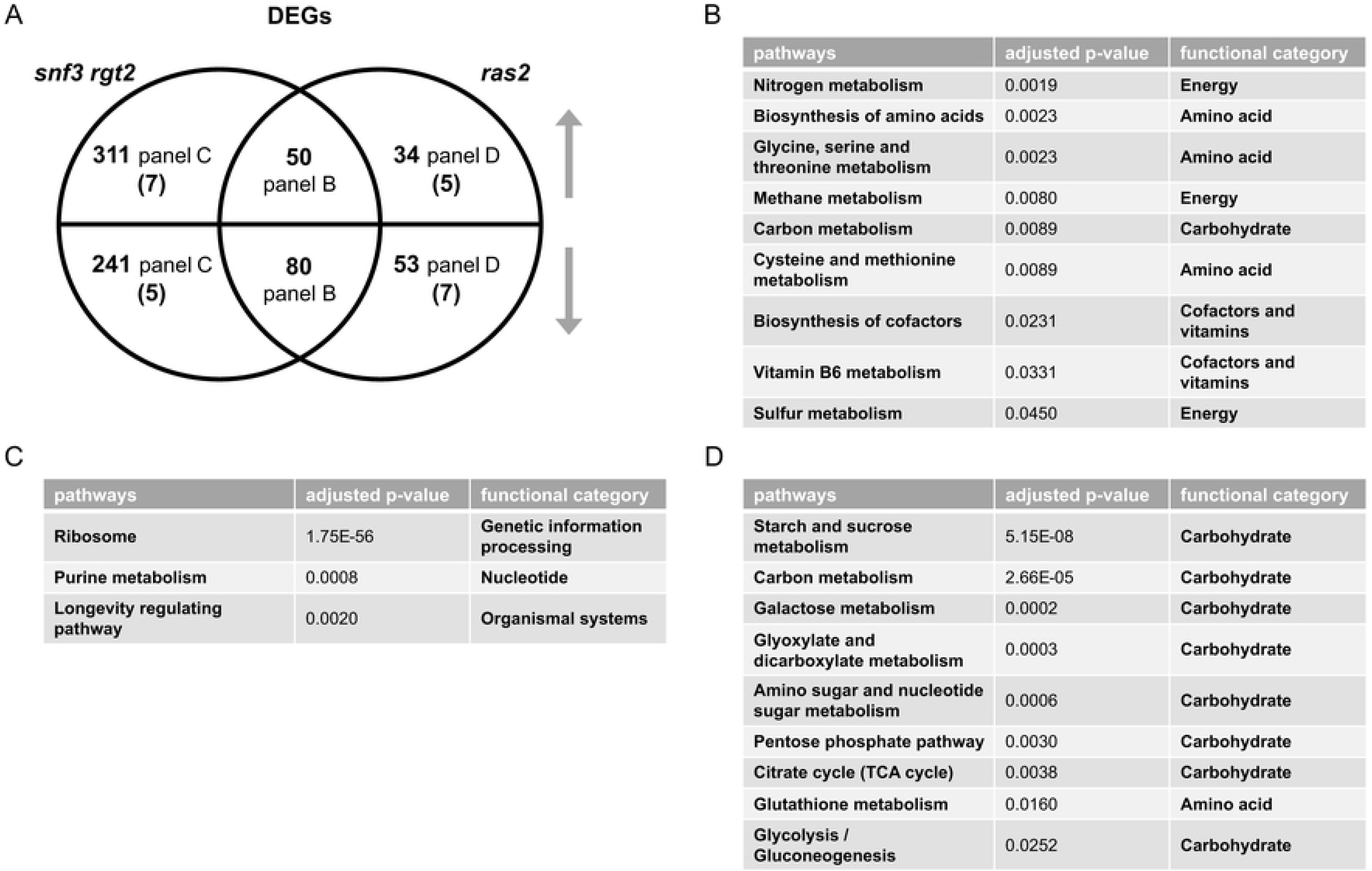
Comparing differentially expressed genes (DEGs) of *snf3 rgt2* and *ras2*. A) Venn diagram of subsets of DEGs, for *snf3 rgt2* vs. wildtype and *ras2* vs. wildtype, after glucose addition to 2%. Upper semicircle shows up-regulated DEGs and lower semicircle shows down-regulated DEGs. Numbers in the overlapping region are shared DEGs regulated in the same direction. Numbers in parenthesis are shared DEGs regulated in the opposite direction, and are placed in the area corresponding to the direction of regulation. DEGs used for ORA analysis that are B) shared and change in the same direction; C) unique to *snf3 rgt2*; D) unique to *ras2.* Listed are all pathways and their functional categories with adjusted p-value <0.05.

**Fig 9.**
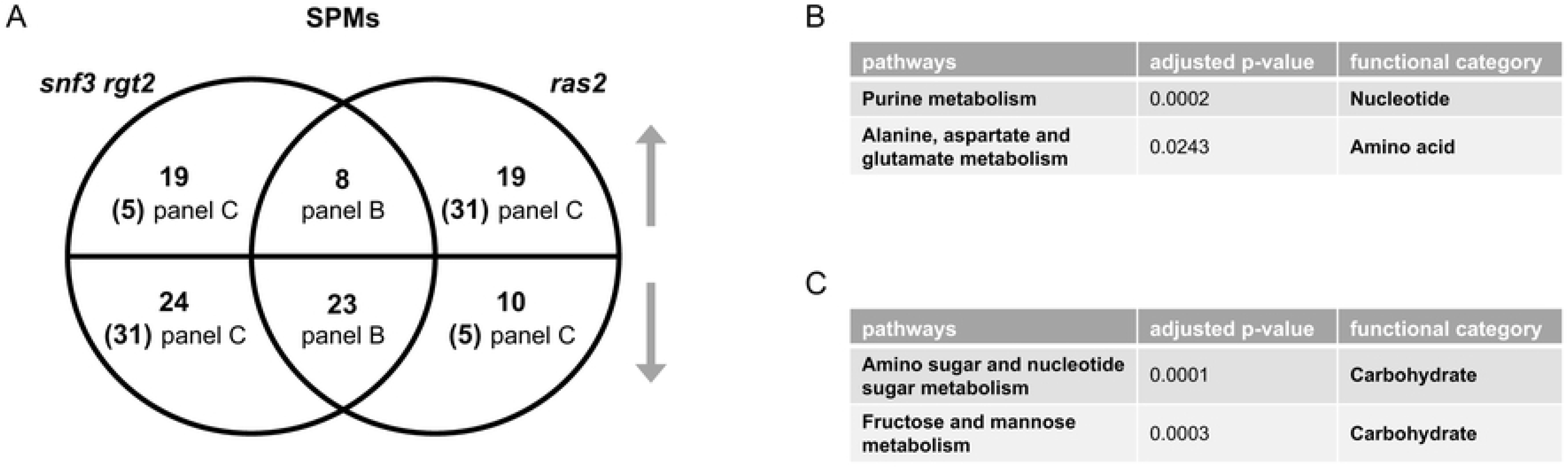
Comparing significantly perturbed metabolites (SPMs) of *snf3 rgt2* and *ras2*. A) Venn diagram of subsets of SPMs, for *snf3 rgt2* vs. wildtype and *ras2* vs. wildtype, after glucose addition. Upper semicircle shows up-regulated SPMs and lower semicircle shows down-regulated SPMs. Numbers in parenthesis are shared SPMs regulated in the opposite direction, and are placed in the area corresponding to the direction of regulation. Numbers in parenthesis are shared SPMs regulated in the opposite direction, and are placed in the area corresponding to the direction of regulation. SPMs used for ORA analysis that are B) shared and change in the same direction; C) shared and change in the opposite direction. Listed are all pathways and their functional categories with adjusted p-value <0.05.

**Fig 10.**
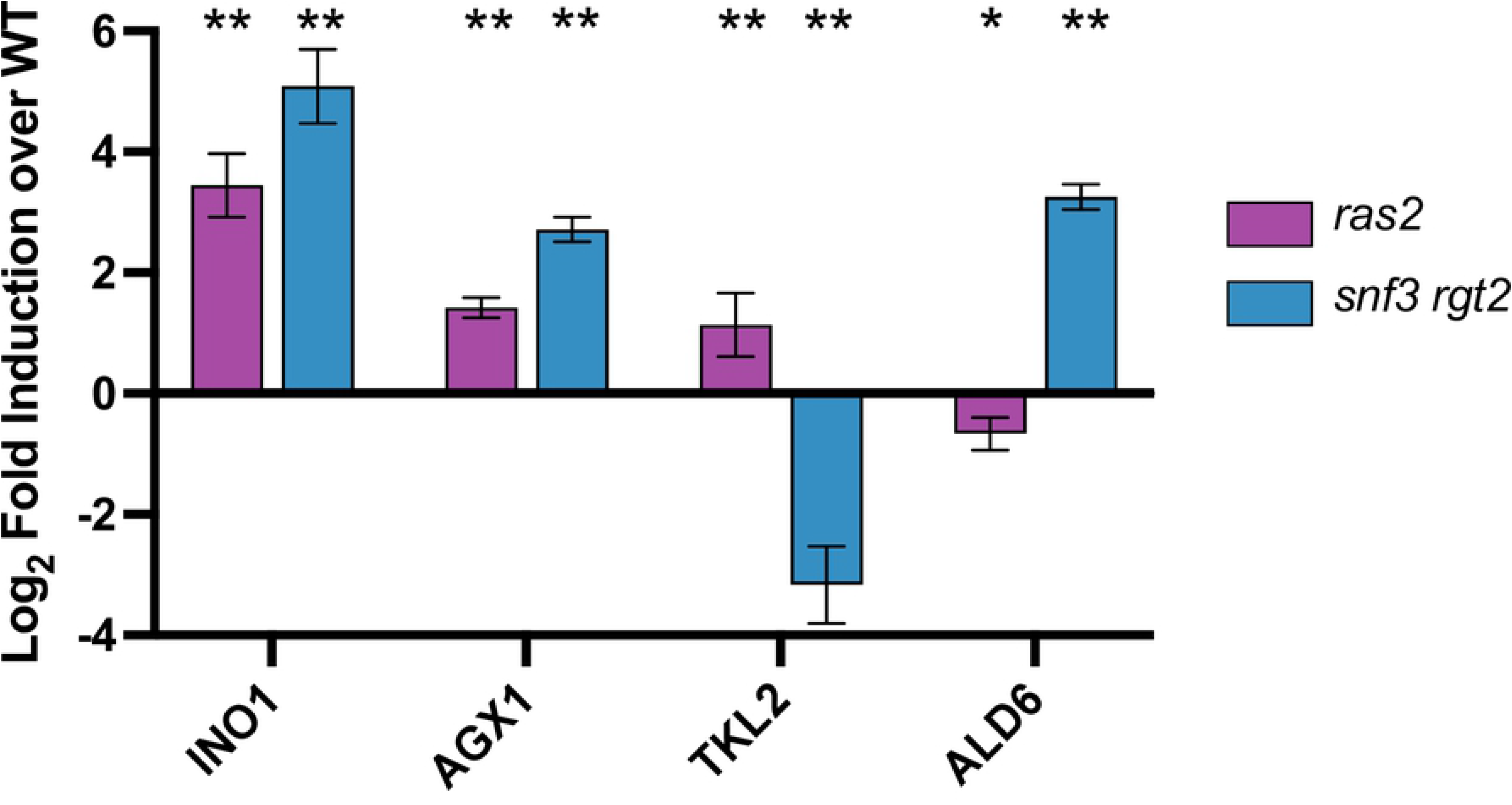
qPCR analysis. Bar plots of qPCR data for *INO1*, *AGX1*, *TKL2*, and ALD6 for *ras2* (purple) and *snf3 rgt2* (blue). X-axis shows target genes; Y-axis shows log_2_ fold induction relative to wildtype. Error bars represent standard error of mean and significance marks are as follows: p<0.01(**), p<0.05(*) as determined via Mann-Whitney U test and adjusted for multiple comparisons with the Benjamini-Hochberg procedure (see Methods).

As noted above, Ras2 and the transceptors had opposing effects on SPMs related to carbohydrate metabolism (Fig 9C). This unexpected effect is further supported by qPCR analysis. As shown in Fig 10, *ras2* and *snf3 rgt2* had significant yet opposing impact on *TKL2*, which encodes an enzyme in the pentose phosphate pathway, and *ALD6*, which produces an aldehyde dehydrogenase involved in pyruvate metabolism. Both enzymes participate directly in carbohydrate metabolism. Therefore, the impact of Snf3 and Rgt2 on carbohydrate metabolism is more limited and antagonistic to that of Ras2. These relationships can be viewed using the KEGG Metabolic Pathways Maps (S2-S5 Figs). Thus, Gpr1 and Ras2 regulate carbohydrate metabolism while Snf3/Rgt2 and Ras2 regulate non-carbohydrate metabolism. We conclude that Ras2 coordinates and integrates signaling by both receptor systems. By integrating transcriptomic and metabolomic measurements, we have taken a major step by identifying new and unexpected functions of Ras2 in the transceptor signaling pathway.

## DISCUSSION

Here we have identified several new and important functions for the glucose sensing apparatus in yeast, comprised of a G protein coupled receptor and a transceptor dimer. Through a systematic analysis of individual gene deletion mutants, we showed how each system contributes - in both shared and unique ways - to transcription and metabolism. In addition, our integration analysis allowed us to confirm and consolidate changes seen at the metabolic or transcriptional level. Whereas the G protein-coupled receptor directs early events in glucose utilization, the transceptors regulate subsequent processes and downstream products of glucose metabolism. While the effects of Ras2 align with those of the G protein coupled receptor, they also align with those of the transceptors. Based on these results, we conclude that Ras2 integrates responses from both receptor systems.

Our approach is distinct from that of prior work on signaling by GPCRs and other cell surface receptors. First, protein components of cell signaling pathways have traditionally been characterized one at a time, often using different readouts for different genes or proteins. Such a piecemeal approach has hindered a comprehensive understanding of the encoded signaling network. Our approach employed comprehensive genome-scale and metabolome-scale (“omic”) measures to quantify differences between mutants lacking individual genes and gene products. Second, our approach was to compare gene deletion mutants in a single-celled organism, one where it is possible to determine functional consequences in the same genetic and epigenetic background, and under identical environmental conditions. By working with yeast we circumvent challenges associated with more complex biological systems, where the structure or topology of the systems is not fully known, the inputs are not static but dynamic (and change over many time scales); such interactions are more likely to be nonlinear and to occur simultaneously at many levels of the biological hierarchy, from molecules to cells to tissues to organs and even to other organisms.

As part of our analysis we compared the function of the individual transceptors, Snf3 and Rgt2, as well as two small G proteins, Ras1 and Ras2. This was done in an effort to determine how these paralogous proteins each contribute to glucose signaling, and with the expectation that such analysis could provide insights into the evolutionary forces that have preserved these gene duplications. Paralogs, or duplicated genes, are especially prevalent in processes related to glucose sensing and utilization in yeast. Apart from the transceptors and Ras proteins, at least four other components of the glucose-sensing pathway (Fig 1) and 8 (out of 12) enzymes responsible for glycolysis [71], are comprised of paralogous gene products. In comparison only 8% of chemical reactions in yeast are executed by paralogs. Systematic deletion of the glycolytic enzymes revealed no defect with respect to gene expression (by microarray), the formation of glycolytic products, or growth rate in a variety of conditions [72]. In keeping with this pattern, our transcriptomics (by RNAseq) and metabolomics analysis (by mass spectrometry) showed that Snf3 and Rgt2 are functionally redundant; that is, deletion of both genes was needed to detect any changes in the thousands of chemical entities measured here (see Data Availability Statement). Of course it is possible that differences in fitness exist but may only be evident under very specific, non-laboratory, growth conditions [71, 73–76].

In the context of previous analysis of gene paralogs, we consider the most significant outcome of our analysis to be that Ras2 (but not Ras1) is required for glucose signaling, and that Ras2 is functionally linked to both receptor systems. Whereas the two receptor systems have distinct roles in signaling, Ras2 appears to integrate the two receptor pathways. Ras2, like Gpr1, directly regulates carbohydrate metabolism. Ras2, like Snf3 and Rgt2, also regulates subsequent processes related to amino acid and nitrogen metabolism. Left unresolved is the role of its paralog Ras1. One possibility is that Ras1 is primarily involved in other aspects of nutrient sensing, as demonstrated for the nitrogen-sensing pathway leading to autophagy [77].

In contrast to Ras2 and Ras1, either Snf3 or Rgt2 can sustain the glucose response. This begs the question, why have both paralogs been retained throughout the course of evolution? Most discussions of gene paralogs have focused on their potential contributions to genetic robustness and phenotypic plasticity [78]. Robustness refers to a number of different mechanisms that stabilize phenotype against genetic or environmental perturbations. An extreme example of robustness is where one of the genes is inactivated and the remaining copy provides enough of the original function to compensate for the loss and ensure survival. In support of this model, several studies in yeast have found that about a third of paralogous gene pairs exhibit negative epistasis [76, 79–81], meaning that deleting both copies produces a significantly larger defect than that of the individual deletions. Robustness could be important when the activity of a duplicated gene product is temporarily disabled in response to changing environmental circumstances, for example through substrate inhibition or feedback phosphorylation. In that case the remaining paralog might compensate for the loss of its sibling by modifying its function through transcriptional reprograming [82, 83], changes in protein stability and abundance [84, 85], or redistribution within the cell [78, 86, 87]. In this way, the overall system may exhibit robustness even while the underlying components exhibit functional plasticity.

Finally, our approach in yeast could guide investigations of functional redundancies in other signaling systems and in other organisms. For example in humans there are three subtypes of Gαi, which assemble with four (out of five) subtypes of Gβ and 12 subtypes of Gγ. Investigators have struggled to find any functional differences among various Gβγ subunit combinations. Another example is the three isoforms of RAS in humans. These proteins were long thought to be functionally interchangeable, since all three share substantial sequence identity in domains responsible for nucleotide binding, GTPase activity, and most effector interactions. However, more recent investigations have shown that HRAS, NRAS and KRAS, when mutated, are each associated with a distinct group of cancer types [88]. An unresolved question is the physiological consequences of these differences with respect to metabolic programming. Moving forward, we believe that the comprehensive, multi-faceted approach taken here could help to provide mechanistic insights to differences among various G proteins in humans.

## METHODS

### Yeast strains

The prototrophic (wildtype) strain used throughout was constructed from BY4741 (MAT**a** *his3*Δ1 *leu2*Δ0 *met15*Δ0 *ura3*Δ0). *HIS3*, *LEU2*, *MET15* and *URA3* were integrated at the endogenous loci with sequence amplified by PCR from S288C strain DNA. All single mutants (*gpr1, gpa2, ras1, ras2, snf3, rgt2*) were constructed by transforming the wildtype strain with corresponding sequence from the Yeast Knock-Out collection that replaces the target gene with KanMX4 [89]. The *snf3 rgt2* double mutant was constructed by switching the mating type of *snf3* from MAT**a** to MATα, with HO expressed from a plasmid, and then mating to an isogenic *rgt2* strain. The diploid was then sporulated and spore products with the *snf3 rgt2* double knock-out were confirmed with PCR.

Cells maintained at 30 °C in Synthetic Complete (SC) (2% glucose) medium were centrifuged and washed twice and then resuspended into 10 mL SC (0.05% glucose) and cultivated for 1 hour. For high and low glucose treatment, 245 μL of 65.5% or 0.05% glucose was added to 10 mL cell culture respectively, each for exactly 2 minutes (metabolomics) or 10 minutes (transcriptomics). Subsequent analysis was performed as described previously [13], and as summarized below.

### Sample preparation for RNA-seq

500 μL of cell culture was centrifuged at 1000 x *g* for 1 minute at 4 °C; the resulting cell pellet was flash frozen by liquid nitrogen. Cells stored at −80 °C were resuspended with 600 μL buffer RLT 1% (v/v) 2-mercaptoethanol from the QIAGEN RNeasy Mini Kit (Cat No.: 74106), transferred to 2 mL OMNI prefilled ceramic bead tubes (SKU: 19-632), loaded onto an OMNI Bead Mill Homogenizer (SKU:19-040E) and agitated three times at 5 m/s for 1 minute at 4 °C while cooled on ice for 3 minutes between each cycle. The resulting lysate was clarified by centrifugation at 11,000 xg and used for total RNA extraction with QIAGEN RNeasy Mini Kit (Cat No.: 74106) with on-column DNase digestion. Extracted total RNA for each sample was evaluated for purity and quantified with the Qubit RNA HS Assay kit (Cat No.: Q32855) and an Invitrogen Qubit 2.0 Fluorometer (Cat No.: Q32866), each according to manufacturer’s instructions.

RNA libraries were prepared with Kapa stranded mRNA-seq kits, with KAPA mRNA Capture Beads (KAPA code: KK8421; Roche Cat No.: 07962207001) through the UNC High Throughput Sequencing Facility. All procedures were according to manufacturer’s instructions.

### RNA Sequence analysis

Quality of raw sequence was checked with the FASTQC algorithm (http://www.bioinformatics.babraham.ac.uk/projects/fastqc/). Sequence alignment to genome indices, generated based on *Saccharomyces cerevisiae* data downloaded from Ensembl.org, was performed with the STAR algorithm [90]. Quantification on the transcriptome level was performed with the SALMON algorithm [91]. Differences in transcript abundance were determined using a negative binomial generalized linear model in DESeq2 package in R [92, 93]. Differentially Expressed Genes (DEGs) were defined as having adjusted p-value <0.05, absolute log2 fold-change >1 and baseMean >100. A series of baseMean thresholds have been tested, including 0, 50 and 100. The conclusion remains unchanged. Therefore, the most stringent threshold (baseMean>100, which filters out >20% of genes) was chosen for data analysis.

PCA analysis was performed using the internal PCA function of DESeq2 package with variance stabilizing transformation (vst) normalized data.

### Transcriptomics pathway enrichment analysis and over-representation analysis

Pathway enrichment analysis for transcriptomics data was performed with ClusterProfiler package in R [63]; Log2 fold-change for each comparison (mutantH vs. wtH) was extracted from corresponding DESeq2 analysis. GSEA analysis was then performed with gseKEGG function, with organism set to ‘sce’ (*Saccharomyces cerevisiae*), permutation number set to 1000, minimal and maximal size for each analyzed geneset as 3 and 200, p-value cutoff set to 0.05, p-value adjustment method set to ‘BH’ (Benjamini-Hochberg).

Over-representation analysis for the corresponding subsection of the Venn diagram was performed with the enrichKEGG function in ClusterProfiler package, with organism set to ‘sce’ (*Saccharomyces cerevisiae*), minimal and maximal size for each analyzed geneset as 3 and 200, p-value cutoff set to 0.05, p-value adjustment method set to ‘BH’ (Benjamini-Hochberg).

### Sample preparation for metabolomics

3 mL of cell culture was mixed with 45 mL cold pure methanol on dry ice and after 5 minutes centrifuged in a precooled rotor (−80 °C). Cell pellets were stored at −80 °C and resuspended with extraction reagent (8:2 methanol-water solution) to 3×10^8^ cell/mL, transferred to 2 mL ceramic bead MagNalyser tubes and subjected to homogenization with Bead Ruptor Elite Bead Mill Homogenizer (OMNI International) at 6.0 m/s for 40 seconds in 2 cycles at room temperature. This homogenization step was repeated twice. After centrifugation at 16,000 xg for 10 minutes at 4 °C, 500 μL of the supernatant was transferred into low-bind 1.7 mL microfuge tubes. Total pools were made by combining an additional 65 μL of the supernatant from each sample and then aliquoting this mixture into low-bind 1.7 mL tubes at a volume of 500 μL. Samples and blanks were dried using a speedvac vacuum concentrator overnight. Following storage at −80 °C, samples were resuspended in 100 μL reconstitution buffer (95:5 water:methanol with 500 ng/mL tryptophan d-5), vortexed at 5000 rpm for 10 minutes, and then centrifuged at room temperature at 16,000 xg for 4 minutes. Supernatant was transferred into autosampler vials for LC-MS.

### UHPLC high-resolution Orbitrap MS metabolomics data acquisition

Metabolomics data were acquired on a Vanquish UHPLC system coupled to a QExactive HF-X Hybrid Quadrupole-Orbitrap Mass Spectrometer (ThermoFisher Scientific, San Jose, CA), as described previously [94]. Our UPLC–MS reversed phase platform was established based on published methods [95, 96]. Metabolites were separated using an HSS T3 C18 column (2.1 × 100 mm, 1.7 μm, Waters Corporation) at 50 °C with binary mobile phase of water (A) and methanol (B), each containing 0.1% formic acid (v/v). The UHPLC linear gradient started from 2% B, and increased to 100% B in 16 minutes, then held for 4 minutes, with the flow rate at 400 μL/minute. The untargeted data were acquired in positive mode from 70 to 1050 m/z using the data-dependent acquisition mode.

### Metabolomics data normalization and filtration

Progenesis QI (version 2.1, Waters Corporation) was used for peak picking, alignment, and normalization as described previously [94]. Samples were randomized and run within two batches with blanks and pools interspersed at a rate of 10%. Starting from the un-normalized data for each of the batch runs, the data were filtered so as to only include signals with an average intensity fold change of 3.0 or greater in the total pools compared to the blanks. Individual samples (including pools, blanks, and study samples) were then normalized to a reference sample that was selected by Progenesis from the total pools via a function named “normalize to all”. Signals were then excluded that were significantly different between pools of batch 1 and pools of batch 2 based on an ANOVA comparison calculated in Progenesis (q <0.05). After normalization and filtration, 2397 signals passed the QC procedures and were used for further analysis.

The filtered and normalized data were mean-centered and Pareto scaled prior to conducting the unsupervised principal component analysis using the ropls R package

### In-house compound identification and annotation

Peaks were identified or annotated by Progenesis QI through matching to an in-house experimental standards library generated by acquiring data for approximately 1000 compounds under conditions identical to study samples, as well as to public databases (including HMDB, METLIN and NIST), as described previously [94]. Identifications and annotations were assigned using available data for retention time (RT), exact mass (MS), MS/MS fragmentation pattern, and isotopic ion pattern. The identification or annotation of each signal is provided in Supporting Information. Signals/metabolites that matched to the in-house experimental standards library by (a) RT, MS, and MS/MS are labeled as OL1, or (b) by RT and MS are labeled OL2a. An OL2b label was provided for signals that match by MS and MS/MS to the in-house library that were outside the retention time tolerance (± 0.5 min) for the standards run under identical conditions. Signals matched to public databases are labeled as PDa (MS and experimental MS/MS), PDb (MS and theoretical MS/MS), PDc (MS and isotopic similarity or adducts), and PDd (MS only) are also provided (Supporting Information).

### Compound annotation, metabolic pathway enrichment analysis and over-representation analysis

Compound annotation and pathway enrichment analysis for metabolomics was performed with the MetaboAnalystR 3.0 package in R [69, 70] (https://www.metaboanalyst.ca/docs/RTutorial.xhtml). For compound annotations, molecular weight tolerance (ppm) was set to 3.0, analytical mode was set to positive and retention time was included. Pathway enrichment analysis was performed with ‘integ’ module (using both Mummichog v2.0 and GSEA) with the yeast KEGG database. The p-value threshold for Mummichog was set at 0.05.

Normalized peak data from Progenesis QI were used as input for MetaboAnalystR. The interaction term estimated how the response amplitude of each mutant is different from wildtype, that is (mutantH-mutantL)-(wtH-wtL). The modeled p-value and t score for the interaction term associated with each peak were then used as inputs for pathway enrichment analysis. Significantly perturbed metabolites (SPMs) were defined as annotations that have adjusted p-value <0.05 (FDR) from the output of MetaboAnalystR. Significantly perturbed pathways were defined as having combined p-value <0.05 (Mummichog and GSEA).

Over-representation analysis for the corresponding subsection of the Venn diagram was performed with the Enrichment Analysis module in MetaboAnalystR, with KEGG ID for each metabolite as the input. FDR adjusted p-value <0.05 was the threshold for over-represented pathways.

### Integration of transcriptomics and metabolomics data

Integration analysis was performed with the ‘joint pathway analysis’ module of MetaboAnalystR (https://www.metaboanalyst.ca/docs/RTutorial.xhtml). Gene input together with log2 fold-change was generated based on the corresponding DESeq2 analysis, with the threshold set as adjusted p-value <0.05, absolute log2 fold-change >1 and baseMean >100 (DEGs); metabolite input together with log2 fold-change was generated based on MetaboAnalystR analysis, with the threshold set as adjusted p-value <0.05 (SPMs). Integration analysis was performed on ‘all pathways’, which includes both metabolic pathways as well as gene-only pathways. Enrichment analysis was performed using ‘Hypergeometric test’. Topology measure was set to ‘Degree Centrality’. Integration method was set to ‘combine queries’, which is a tight integration method with genes and metabolites pooled into a single query and used to perform enrichment analysis within their “pooled universe”. Significantly enriched pathways were defined as having FDR adjusted p-value <0.05.

### Yeast RNA extraction, DNase treatment, and reverse transcription for qPCR

RNA was extracted from cells using hot acid phenol. TES solution (10 mM Tris-HCl, pH 7.5; 10 mM EDTA; 0.5% SDS) was used to resuspend pellets then the resuspension was incubated for one hour at 65°C. The RNA was separated via phenol-chloroform extraction and any residual DNA was degraded with RQ1 DNase (Promega). To further purify the RNA, RNeasy mini kit (Qiagen) was used and the final RNA concentration was determined via spectrophotometry with a NanoDrop One (ThermoFisher Scientific). cDNA was produced via reverse transcription from 250 ng RNA using a High-Capacity cDNA Reverse Transcription Kit (ThermoFisher Scientific) following manufacturer’s protocol.

### qPCR

qPCR primers were ordered from Integrated DNA Technologies:

YER100W_FWD primer: 5’ GAAGCCACGACAGGATCAAT 3’

YER100W_REV primer: 5’ ATCCCCCTCATCCAATTTTC 3’

YBR117C_FWD: 5’ GTCACTCATGCGCTCTTCTG 3’

YBR117C_REV: 5’ GAGTCGGAAATGGGAAAGCC 3’

YPL061W_FWD: 5’ GGCGCCAAGATCTTAACTGG 3’

YPL061W_REV: 5’ CCACCTTCAAACCTGTGCTC 3’

YJL153C_FWD: 5’ CATGGTTAGCCCAAACGACT 3’

YJL153C_REV: 5’ CGTGGTTACGTTGCCTTTTT 3’

YFL030W_FWD: 5’ TGATCCCAGGCCCCATTATC 3’

YFL030W_REV: 5’ AATATGTCCCACCCCAACGT 3’

To perform qPCR, cDNA was diluted 50-fold and amplified with SsoAdvanced Universal SYBR Green Supermix (Bio-Rad) following manufacturer’s protocols with adjustments: 45 cycles were used to increase amplification and anneal/extension time was extended to 45 seconds. qPCR was performed in technical triplicate for each of the six biological replicates per genotype. CFX Maestro Software (Bio-Rad) was used to determine the threshold cycle (C_t_). ΔC_t_ values were determined in reference to YER100W and final ΔΔC_t_ values were calculated and normalized in reference to wildtype cells. p values were calculated on ΔC_t_ values between genotypes via independent, non-parametric, one-tailed Mann-Whitney U tests with the expected change in expression as was found by RNAseq. One exception was that of *TKL2* in *snf3 rgt2 vs*. wildtype comparison in which the RNAseq data did not yield a statistically significant change, in this case a two-sided Mann-Whitney U test was applied. The Benjamini-Hochberg Procedure was used to correct for multiple comparisons.

## SUPPORTING INFORMATION

**S1 Fig. PCA plots.** For A) transcriptomics and B) metabolomics, X-axis shows PC1 with the percentage of explained variance and Y-axis shows PC2 with the percentage of explained variance. Data are scaled as detailed in Methods. Wildtype (black), *gpr1* (red), *gpa2* (orange), *snf3 rgt2* (blue), *ras1* (green), *ras2* (purple). Low glucose (L, 0.05% glucose)-triangles, high glucose (H, 2% glucose)-circles.

**S2 Fig. Overview of DEGs and SPMs regulated by *GPR1* projected on KEGG Metabolic Pathway.** The map is color coded to delineate carbohydrate metabolism (blue), glycan biosynthesis and metabolism (cyan), amino acid metabolism (yellow), nucleotide metabolism (red), lipid metabolism (teal), metabolism of cofactors and vitamins (pink). Highlighted are the DEGs (black lines) and SPMs (black dots) for *gpr1* integration analysis and gray boxes are used to delineate clusters associated with a specific pathway.

**S3 Fig. Overview of DEGs and SPMs regulated by *SNF3* and *RGT2* projected on KEGG Metabolic Pathway.** The map is color coded as in S2 Fig. Highlighted are the DEGs (black lines) and SPMs (black dots) for *snf3 rgt2* integration analysis and gray boxes are used to delineate clusters associated with a specific pathway.

**S4 Fig. Overview of DEGs and SPMs regulated by *GPA2* projected on KEGG Metabolic Pathway.** The map is color coded as in S2 Fig. Highlighted are the DEGs (black lines) and SPMs (black dots) for *gpa2* integration analysis and gray boxes are used to delineate clusters associated with a specific pathway.

**S5 Fig. Overview of DEGs and SPMs regulated by *RAS2* projected on KEGG Metabolic Pathway.** The map is color coded as in S2 Fig. Highlighted are the DEGs (black lines) and SPMs (black dots) for *ras2* integration analysis and gray boxes are used to delineate clusters associated with a specific pathway.

**S1 Table. Single-omics analysis results for wildtype between 2% (H) and 0.05% (L) glucose.** First block shows GSEA for transcriptomics with adjusted p-value <0.05, arranged in ascending order; second block shows MetaboAnalystR pathway enrichment analysis for metabolomics with combined p-value <0.05 arranged in ascending order. Reproduced from [13].

**S2 Table.** Results and statistics of transcriptomics, metabolomics and multi -omics integration for *gpr1*, each as a separate sheet.

**S3 Table.** Results and statistics of transcriptomics, metabolomics and multi -omics integration for *snf3 rgt2*, each as a separate sheet.

**S4 Table.** List of DEGs for each subset of the Venn diagram in Fig 2A.

**S5 Table.** List of SPMs for each subset of the Venn diagram in Fig 3A.

**S6 Table.** In house compound identification.

**S7 Table.** Results and statistics of transcriptomics, metabolomics and multi -omics integration for *gpa2*, each as a separate sheet.

**S8 Table.** Results and statistics of transcriptomics, metabolomics and multi -omics integration for *ras2*, each as a separate sheet.

**S9 Table.** List of DEGs for each subset of the Venn diagram in Fig 5A.

**S10 Table.** List of SPMs for each subset of the Venn diagram in Fig 6A.

**S11 Table.** List of DEGs for each subset of the Venn diagram in Fig 8A.

**S12 Table.** List of SPMs for each subset of the Venn diagram in Fig 9A

## REFERENCES

1. Kresnowati MT, van Winden WA, Almering MJ, ten Pierick A, Ras C, Knijnenburg TA, et al. When transcriptome meets metabolome: fast cellular responses of yeast to sudden relief of glucose limitation. Mol Syst Biol. 2006;2:49. Epub 2006/09/14. doi: 10.1038/msb4100083. PubMed PMID: 16969341; PubMed Central PMCID: PMCPMC1681515.

2. Castrillo JI, Zeef LA, Hoyle DC, Zhang N, Hayes A, Gardner DC, et al. Growth control of the eukaryote cell: a systems biology study in yeast. J Biol. 2007;6(2):4. Epub 2007/04/19. doi: 10.1186/jbiol54. PubMed PMID: 17439666; PubMed Central PMCID: PMCPMC2373899.

3. Gutteridge A, Pir P, Castrillo JI, Charles PD, Lilley KS, Oliver SG. Nutrient control of eukaryote cell growth: a systems biology study in yeast. BMC Biol. 2010;8:68. Epub 2010/05/26. doi: 10.1186/1741-7007-8-68. PubMed PMID: 20497545; PubMed Central PMCID: PMCPMC2895586.

4. Dikicioglu D, Karabekmez E, Rash B, Pir P, Kirdar B, Oliver SG. How yeast re-programmes its transcriptional profile in response to different nutrient impulses. BMC Syst Biol. 2011;5:148. Epub 2011/09/29. doi: 10.1186/1752-0509-5-148. PubMed PMID: 21943358; PubMed Central PMCID: PMCPMC3224505.

5. Yun CW, Tamaki H, Nakayama R, Yamamoto K, Kumagai H. G-protein coupled receptor from yeast Saccharomyces cerevisiae. Biochem Biophys Res Commun. 1997;240(2):287–92.

6. Yun CW, Tamaki H, Nakayama R, Yamamoto K, Kumagai H. Gpr1p, a putative G-protein coupled receptor, regulates glucose-dependent cellular cAMP level in yeast Saccharomyces cerevisiae. Biochem Biophys Res Commun. 1998;252(1):29–33.

7. Xue Y, Batlle M, Hirsch JP. GPR1 encodes a putative G protein-coupled receptor that associates with the Gpa2p Galpha subunit and functions in a Ras-independent pathway. Embo J. 1998;17(7):1996–2007.

8. Colombo S, Ma P, Cauwenberg L, Winderickx J, Crauwels M, Teunissen A, et al. Involvement of distinct G-proteins, Gpa2 and Ras, in glucose- and intracellular acidification-induced cAMP signalling in the yeast Saccharomyces cerevisiae. Embo J. 1998;17(12):3326–41.

9. Kraakman L, Lemaire K, Ma P, Teunissen AW, Donaton MC, Van Dijck P, et al. A Saccharomyces cerevisiae G-protein coupled receptor, Gpr1, is specifically required for glucose activation of the cAMP pathway during the transition to growth on glucose. Mol Microbiol. 1999;32(5):1002–12.

10. Lorenz MC, Pan X, Harashima T, Cardenas ME, Xue Y, Hirsch JP, et al. The G protein-coupled receptor gpr1 is a nutrient sensor that regulates pseudohyphal differentiation in Saccharomyces cerevisiae. Genetics. 2000;154(2):609–22.

11. Lemaire K, Van de Velde S, Van Dijck P, Thevelein JM. Glucose and sucrose act as agonist and mannose as antagonist ligands of the G protein-coupled receptor Gpr1 in the yeast Saccharomyces cerevisiae. Mol Cell. 2004;16(2):293–9. PubMed PMID: 15494315.

12. Zeller CE, Parnell SC, Dohlman HG. The RACK1 ortholog Asc1 functions as a G-protein beta subunit coupled to glucose responsiveness in yeast. J Biol Chem. 2007;282(34):25168–76. PubMed PMID: 17591772.

13. Li S, Li Y, Rushing BR, Harris SE, McRitchie SL, Jones JC, et al. Multi-omics analysis of glucose-mediated signaling by a moonlighting Gbeta protein Asc1/RACK1. PLoS Genet. 2021;17(7):e1009640. Epub 2021/07/03. doi: 10.1371/journal.pgen.1009640. PubMed PMID: 34214075; PubMed Central PMCID: PMCPMC8282090 paid employee of Metabolon, a for-profit company. Metabolon provided no data, data analysis, employment or consultancy, and claims no rights to possible patents or products that may arise from the research.

14. Broek D, Toda T, Michaeli T, Levin L, Birchmeier C, Zoller M, et al. The S. cerevisiae CDC25 gene product regulates the RAS/adenylate cyclase pathway. Cell. 1987;48(5):789–99. PubMed PMID: 3545497.

15. Munder T, Kuntzel H. Glucose-induced cAMP signaling in Saccharomyces cerevisiae is mediated by the CDC25 protein. FEBS Lett. 1989;242(2):341–5. PubMed PMID: 2536619.

16. Crechet JB, Poullet P, Mistou MY, Parmeggiani A, Camonis J, Boy-Marcotte E, et al. Enhancement of the GDP-GTP exchange of RAS proteins by the carboxyl-terminal domain of SCD25. Science. 1990;248(4957):866–8. PubMed PMID: 2188363.

17. Jones S, Vignais ML, Broach JR. The CDC25 protein of Saccharomyces cerevisiae promotes exchange of guanine nucleotides bound to ras. Mol Cell Biol. 1991;11(5):2641–6. Epub 1991/05/01. PubMed PMID: 2017169; PubMed Central PMCID: PMC360033.

18. Papasavvas S, Arkinstall S, Reid J, Payton M. Yeast alpha-mating factor receptor and G-protein-linked adenylyl cyclase inhibition requires RAS2 and GPA2 activities. Biochem Biophys Res Commun. 1992;184(3):1378–85. PubMed PMID: 1317171.

19. Boy-Marcotte E, Ikonomi P, Jacquet M. SDC25, a dispensable Ras guanine nucleotide exchange factor of Saccharomyces cerevisiae differs from CDC25 by its regulation. Mol Biol Cell. 1996;7(4):529–39. PubMed PMID: 8730097; PubMed Central PMCID: PMC275907.

20. Gross A, Winograd S, Marbach I, Levitzki A. The N-terminal half of Cdc25 is essential for processing glucose signaling in Saccharomyces cerevisiae. Biochemistry. 1999;38(40):13252–62. PubMed PMID: 10529198.

21. VanderSluis B, Hess DC, Pesyna C, Krumholz EW, Syed T, Szappanos B, et al. Broad metabolic sensitivity profiling of a prototrophic yeast deletion collection. Genome Biol. 2014;15(4):R64. Epub 2014/04/12. doi: 10.1186/gb-2014-15-4-r64. PubMed PMID: 24721214; PubMed Central PMCID: PMCPMC4053978.

22. Powers S, Kataoka T, Fasano O, Goldfarb M, Strathern J, Broach J, et al. Genes in S. cerevisiae encoding proteins with domains homologous to the mammalian ras proteins. Cell. 1984;36(3):607–12. PubMed PMID: 6365329.

23. Toda T, Uno I, Ishikawa T, Powers S, Kataoka T, Broek D, et al. In yeast, RAS proteins are controlling elements of adenylate cyclase. Cell. 1985;40(1):27–36. Epub 1985/01/01. doi: 0092-8674(85)90305-8 [pii]. PubMed PMID: 2981630.

24. Uno I, Mitsuzawa H, Matsumoto K, Tanaka K, Oshima T, Ishikawa T. Reconstitution of the GTP-dependent adenylate cyclase from products of the yeast CYR1 and RAS2 genes in Escherichia coli. Proc Natl Acad Sci U S A. 1985;82(23):7855–9. Epub 1985/12/01. PubMed PMID: 2999779; PubMed Central PMCID: PMC390868.

25. Field J, Nikawa J, Broek D, MacDonald B, Rodgers L, Wilson IA, et al. Purification of a RAS-responsive adenylyl cyclase complex from Saccharomyces cerevisiae by use of an epitope addition method. Mol Cell Biol. 1988;8(5):2159–65. PubMed PMID: 2455217; PubMed Central PMCID: PMC363397.

26. Nakafuku M, Obara T, Kaibuchi K, Miyajima I, Miyajima A, Itoh H, et al. Isolation of a second yeast Saccharomyces cerevisiae gene (GPA2) coding for guanine nucleotide-binding regulatory protein: studies on its structure and possible functions. Proc Natl Acad Sci U S A. 1988;85(5):1374–8.

27. Field J, Xu HP, Michaeli T, Ballester R, Sass P, Wigler M, et al. Mutations of the adenylyl cyclase gene that block RAS function in Saccharomyces cerevisiae. Science. 1990;247(4941):464–7. PubMed PMID: 2405488.

28. Suzuki N, Choe HR, Nishida Y, Yamawaki-Kataoka Y, Ohnishi S, Tamaoki T, et al. Leucine-rich repeats and carboxyl terminus are required for interaction of yeast adenylate cyclase with RAS proteins. Proc Natl Acad Sci U S A. 1990;87(22):8711–5. PubMed PMID: 2247439; PubMed Central PMCID: PMCPMC55029.

29. Mintzer KA, Field J. Interactions between adenylyl cyclase, CAP and RAS from Saccharomyces cerevisiae. Cell Signal. 1994;6(6):681–94. PubMed PMID: 7531994.

30. Bhattacharya S, Chen L, Broach JR, Powers S. Ras membrane targeting is essential for glucose signaling but not for viability in yeast. Proc Natl Acad Sci U S A. 1995;92(7):2984–8.

31. Kubler E, Mosch HU, Rupp S, Lisanti MP. Gpa2p, a G-protein alpha-subunit, regulates growth and pseudohyphal development in Saccharomyces cerevisiae via a cAMP-dependent mechanism. J Biol Chem. 1997;272(33):20321–3.

32. Rolland F, De Winde JH, Lemaire K, Boles E, Thevelein JM, Winderickx J. Glucose-induced cAMP signalling in yeast requires both a G-protein coupled receptor system for extracellular glucose detection and a separable hexose kinase-dependent sensing process. Mol Microbiol. 2000;38(2):348–58. PubMed PMID: 11069660.

33. Wang Y, Pierce M, Schneper L, Guldal CG, Zhang X, Tavazoie S, et al. Ras and Gpa2 mediate one branch of a redundant glucose signaling pathway in yeast. PLoS Biol. 2004;2(5):E128. doi: 10.1371/journal.pbio.0020128. PubMed PMID: 15138498; PubMed Central PMCID: PMC406390.

34. Matsumoto K, Uno I, Oshima Y, Ishikawa T. Isolation and characterization of yeast mutants deficient in adenylate cyclase and cAMP-dependent protein kinase. Proc Natl Acad Sci U S A. 1982;79(7):2355–9. PubMed PMID: 6285379; PubMed Central PMCID: PMC346192.

35. Kataoka T, Broek D, Wigler M. DNA sequence and characterization of the S. cerevisiae gene encoding adenylate cyclase. Cell. 1985;43(2 Pt 1):493–505. PubMed PMID: 2934138.

36. Casperson GF, Walker N, Bourne HR. Isolation of the gene encoding adenylate cyclase in Saccharomyces cerevisiae. Proc Natl Acad Sci U S A. 1985;82(15):5060–3. PubMed PMID: 2991907; PubMed Central PMCID: PMC390498.

37. Harashima T, Heitman J. The Galpha protein Gpa2 controls yeast differentiation by interacting with kelch repeat proteins that mimic Gbeta subunits. Mol Cell. 2002;10(1):163–73. PubMed PMID: 12150916.

38. Toda T, Cameron S, Sass P, Zoller M, Scott JD, McMullen B, et al. Cloning and characterization of BCY1, a locus encoding a regulatory subunit of the cyclic AMP-dependent protein kinase in Saccharomyces cerevisiae. Mol Cell Biol. 1987;7(4):1371–7. Epub 1987/04/01. PubMed PMID: 3037314; PubMed Central PMCID: PMC365223.

39. Toda T, Cameron S, Sass P, Zoller M, Wigler M. Three different genes in S. cerevisiae encode the catalytic subunits of the cAMP-dependent protein kinase. Cell. 1987;50(2):277–87. Epub 1987/07/17. doi: 0092-8674(87)90223-6 [pii]. PubMed PMID: 3036373.

40. Cannon JF, Tatchell K. Characterization of Saccharomyces cerevisiae genes encoding subunits of cyclic AMP-dependent protein kinase. Mol Cell Biol. 1987;7(8):2653–63. PubMed PMID: 2823100; PubMed Central PMCID: PMC367881.

41. Robertson LS, Fink GR. The three yeast A kinases have specific signaling functions in pseudohyphal growth. Proc Natl Acad Sci U S A. 1998;95(23):13783–7.

42. Pan X, Heitman J. Cyclic AMP-dependent protein kinase regulates pseudohyphal differentiation in Saccharomyces cerevisiae. Mol Cell Biol. 1999;19(7):4874–87.

43. Robertson LS, Causton HC, Young RA, Fink GR. The yeast A kinases differentially regulate iron uptake and respiratory function. Proc Natl Acad Sci U S A. 2000;97(11):5984–8. doi: 10.1073/pnas.100113397. PubMed PMID: 10811893; PubMed Central PMCID: PMC18545.

44. Ptacek J, Devgan G, Michaud G, Zhu H, Zhu X, Fasolo J, et al. Global analysis of protein phosphorylation in yeast. Nature. 2005;438(7068):679–84. Epub 2005/12/02. doi: nature04187 [pii] 10.1038/nature04187. PubMed PMID: 16319894.

45. Neigeborn L, Schwartzberg P, Reid R, Carlson M. Null mutations in the SNF3 gene of Saccharomyces cerevisiae cause a different phenotype than do previously isolated missense mutations. Mol Cell Biol. 1986;6(11):3569–74. PubMed PMID: 3540596; PubMed Central PMCID: PMC367116.

46. Ozcan S, Dover J, Rosenwald AG, Wolfl S, Johnston M. Two glucose transporters in Saccharomyces cerevisiae are glucose sensors that generate a signal for induction of gene expression. Proc Natl Acad Sci U S A. 1996;93(22):12428–32. PubMed PMID: 8901598; PubMed Central PMCID: PMC38008.

47. Ozcan S, Dover J, Johnston M. Glucose sensing and signaling by two glucose receptors in the yeast Saccharomyces cerevisiae. EMBO J. 1998;17(9):2566–73. doi: 10.1093/emboj/17.9.2566. PubMed PMID: 9564039; PubMed Central PMCID: PMC1170598.

48. Schmidt MC, McCartney RR, Zhang X, Tillman TS, Solimeo H, Wolfl S, et al. Std1 and Mth1 proteins interact with the glucose sensors to control glucose-regulated gene expression in Saccharomyces cerevisiae. Mol Cell Biol. 1999;19(7):4561–71. PubMed PMID: 10373505; PubMed Central PMCID: PMC84254.

49. Lafuente MJ, Gancedo C, Jauniaux JC, Gancedo JM. Mth1 receives the signal given by the glucose sensors Snf3 and Rgt2 in Saccharomyces cerevisiae. Mol Microbiol. 2000;35(1):161–72. PubMed PMID: 10632886.

50. Spielewoy N, Flick K, Kalashnikova TI, Walker JR, Wittenberg C. Regulation and recognition of SCFGrr1 targets in the glucose and amino acid signaling pathways. Mol Cell Biol. 2004;24(20):8994–9005. doi: 10.1128/MCB.24.20.8994-9005.2004. PubMed PMID: 15456873; PubMed Central PMCID: PMC517892.

51. Moriya H, Johnston M. Glucose sensing and signaling in Saccharomyces cerevisiae through the Rgt2 glucose sensor and casein kinase I. Proc Natl Acad Sci U S A. 2004;101(6):1572–7. doi: 10.1073/pnas.0305901101. PubMed PMID: 14755054; PubMed Central PMCID: PMC341776.

52. Pasula S, Jouandot D, 2nd, Kim JH. Biochemical evidence for glucose-independent induction of HXT expression in Saccharomyces cerevisiae. FEBS Lett. 2007;581(17):3230–4. doi: 10.1016/j.febslet.2007.06.013. PubMed PMID: 17586499; PubMed Central PMCID: PMC2040036.

53. Tomas-Cobos L, Sanz P. Active Snf1 protein kinase inhibits expression of the Saccharomyces cerevisiae HXT1 glucose transporter gene. Biochem J. 2002;368(Pt 2):657–63. doi: 10.1042/BJ20020984. PubMed PMID: 12220226; PubMed Central PMCID: PMC1223017.

54. Kim JH, Polish J, Johnston M. Specificity and regulation of DNA binding by the yeast glucose transporter gene repressor Rgt1. Mol Cell Biol. 2003;23(15):5208–16. PubMed PMID: 12861007; PubMed Central PMCID: PMC165726.

55. Flick KM, Spielewoy N, Kalashnikova TI, Guaderrama M, Zhu Q, Chang HC, et al. Grr1-dependent inactivation of Mth1 mediates glucose-induced dissociation of Rgt1 from HXT gene promoters. Mol Biol Cell. 2003;14(8):3230–41. doi: 10.1091/mbc.E03-03-0135. PubMed PMID: 12925759; PubMed Central PMCID: PMC181563.

56. Mosley AL, Lakshmanan J, Aryal BK, Ozcan S. Glucose-mediated phosphorylation converts the transcription factor Rgt1 from a repressor to an activator. J Biol Chem. 2003;278(12):10322–7. doi: 10.1074/jbc.M212802200. PubMed PMID: 12527758.

57. Lakshmanan J, Mosley AL, Ozcan S. Repression of transcription by Rgt1 in the absence of glucose requires Std1 and Mth1. Curr Genet. 2003;44(1):19–25. doi: 10.1007/s00294-003-0423-2. PubMed PMID: 14508605.

58. Polish JA, Kim JH, Johnston M. How the Rgt1 transcription factor of Saccharomyces cerevisiae is regulated by glucose. Genetics. 2005;169(2):583–94. doi: 10.1534/genetics.104.034512. PubMed PMID: 15489524; PubMed Central PMCID: PMC1449106.

59. Kim JH, Johnston M. Two glucose-sensing pathways converge on Rgt1 to regulate expression of glucose transporter genes in Saccharomyces cerevisiae. J Biol Chem. 2006;281(36):26144–9. doi: 10.1074/jbc.M603636200. PubMed PMID: 16844691.

60. Palomino A, Herrero P, Moreno F. Tpk3 and Snf1 protein kinases regulate Rgt1 association with Saccharomyces cerevisiae HXK2 promoter. Nucleic Acids Res. 2006;34(5):1427–38. doi: 10.1093/nar/gkl028. PubMed PMID: 16528100; PubMed Central PMCID: PMC1401511.

61. Jouandot D, 2nd, Roy A, Kim JH. Functional dissection of the glucose signaling pathways that regulate the yeast glucose transporter gene (HXT) repressor Rgt1. J Cell Biochem. 2011;112(11):3268–75. doi: 10.1002/jcb.23253. PubMed PMID: 21748783; PubMed Central PMCID: PMC3341738.

62. Roy A, Shin YJ, Cho KH, Kim JH. Mth1 regulates the interaction between the Rgt1 repressor and the Ssn6-Tup1 corepressor complex by modulating PKA-dependent phosphorylation of Rgt1. Mol Biol Cell. 2013;24(9):1493–503. doi: 10.1091/mbc.E13-01-0047. PubMed PMID: 23468525; PubMed Central PMCID: PMC3639059.

63. Yu G, Wang LG, Han Y, He QY. clusterProfiler: an R package for comparing biological themes among gene clusters. OMICS. 2012;16(5):284–7. Epub 2012/03/30. doi: 10.1089/omi.2011.0118. PubMed PMID: 22455463; PubMed Central PMCID: PMCPMC3339379.

64. Kanehisa M, Goto S. KEGG: kyoto encyclopedia of genes and genomes. Nucleic Acids Res. 2000;28(1):27–30. Epub 1999/12/11. doi: 10.1093/nar/28.1.27. PubMed PMID: 10592173; PubMed Central PMCID: PMCPMC102409.

65. Kanehisa M. Toward understanding the origin and evolution of cellular organisms. Protein Sci. 2019;28(11):1947–51. Epub 2019/08/24. doi: 10.1002/pro.3715. PubMed PMID: 31441146; PubMed Central PMCID: PMCPMC6798127.

66. Kanehisa M, Furumichi M, Sato Y, Ishiguro-Watanabe M, Tanabe M. KEGG: integrating viruses and cellular organisms. Nucleic Acids Res. 2020. Epub 2020/10/31. doi: 10.1093/nar/gkaa970. PubMed PMID: 33125081.

67. Zhang N, Cao L. Starvation signals in yeast are integrated to coordinate metabolic reprogramming and stress response to ensure longevity. Curr Genet. 2017;63(5):839–43. Epub 2017/04/27. doi: 10.1007/s00294-017-0697-4. PubMed PMID: 28444510; PubMed Central PMCID: PMCPMC5605593.

68. Li S, Park Y, Duraisingham S, Strobel FH, Khan N, Soltow QA, et al. Predicting network activity from high throughput metabolomics. PLoS Comput Biol. 2013;9(7):e1003123. Epub 2013/07/19. doi: 10.1371/journal.pcbi.1003123. PubMed PMID: 23861661; PubMed Central PMCID: PMCPMC3701697.

69. Chong J, Wishart DS, Xia J. Using MetaboAnalyst 4.0 for Comprehensive and Integrative Metabolomics Data Analysis. Curr Protoc Bioinformatics. 2019;68(1):e86. Epub 2019/11/23. doi: 10.1002/cpbi.86. PubMed PMID: 31756036.

70. Pang Z, Chong J, Li S, Xia J. MetaboAnalystR 3.0: Toward an Optimized Workflow for Global Metabolomics. Metabolites. 2020;10(5). Epub 2020/05/13. doi: 10.3390/metabo10050186. PubMed PMID: 32392884.

71. Bradley PH, Gibney PA, Botstein D, Troyanskaya OG, Rabinowitz JD. Minor Isozymes Tailor Yeast Metabolism to Carbon Availability. mSystems. 2019;4(1). Epub 2019/03/06. doi: 10.1128/mSystems.00170-18. PubMed PMID: 30834327; PubMed Central PMCID: PMCPMC6392091.

72. Solis-Escalante D, Kuijpers NG, Barrajon-Simancas N, van den Broek M, Pronk JT, Daran JM, et al. A Minimal Set of Glycolytic Genes Reveals Strong Redundancies in Saccharomyces cerevisiae Central Metabolism. Eukaryot Cell. 2015;14(8):804–16. Epub 2015/06/14. doi: 10.1128/EC.00064-15. PubMed PMID: 26071034; PubMed Central PMCID: PMCPMC4519752.

73. Giaever G, Chu AM, Ni L, Connelly C, Riles L, Veronneau S, et al. Functional profiling of the Saccharomyces cerevisiae genome. Nature. 2002;418(6896):387–91.

74. Papp B, Pal C, Hurst LD. Metabolic network analysis of the causes and evolution of enzyme dispensability in yeast. Nature. 2004;429(6992):661–4. Epub 2004/06/11. doi: 10.1038/nature02636. PubMed PMID: 15190353.

75. Ihmels J, Collins SR, Schuldiner M, Krogan NJ, Weissman JS. Backup without redundancy: genetic interactions reveal the cost of duplicate gene loss. Mol Syst Biol. 2007;3:86. Epub 2007/03/29. doi: 10.1038/msb4100127. PubMed PMID: 17389874; PubMed Central PMCID: PMCPMC1847942.

76. DeLuna A, Vetsigian K, Shoresh N, Hegreness M, Colon-Gonzalez M, Chao S, et al. Exposing the fitness contribution of duplicated genes. Nat Genet. 2008;40(5):676–81. Epub 2008/04/15. doi: 10.1038/ng.123. PubMed PMID: 18408719.

77. Jin X, Starke S, Li Y, Sethupathi S, Kung G, Dodhiawala P, et al. Nitrogen Starvation-induced Phosphorylation of Ras1 Protein and Its Potential Role in Nutrient Signaling and Stress Response. J Biol Chem. 2016;291(31):16231–9. Epub 2016/06/05. doi: 10.1074/jbc.M115.713206. PubMed PMID: 27261458; PubMed Central PMCID: PMCPMC4965571.

78. Nijhout HF, Sadre-Marandi F, Best J, Reed MC. Systems Biology of Phenotypic Robustness and Plasticity. Integr Comp Biol. 2017;57(2):171–84. Epub 2017/09/02. doi: 10.1093/icb/icx076. PubMed PMID: 28859407.

79. Dean EJ, Davis JC, Davis RW, Petrov DA. Pervasive and persistent redundancy among duplicated genes in yeast. PLoS Genet. 2008;4(7):e1000113. Epub 2008/07/08. doi: 10.1371/journal.pgen.1000113. PubMed PMID: 18604285; PubMed Central PMCID: PMCPMC2440806.

80. Musso G, Costanzo M, Huangfu M, Smith AM, Paw J, San Luis BJ, et al. The extensive and condition-dependent nature of epistasis among whole-genome duplicates in yeast. Genome Res. 2008;18(7):1092–9. Epub 2008/05/09. doi: 10.1101/gr.076174.108. PubMed PMID: 18463300; PubMed Central PMCID: PMCPMC2493398.

81. VanderSluis B, Bellay J, Musso G, Costanzo M, Papp B, Vizeacoumar FJ, et al. Genetic interactions reveal the evolutionary trajectories of duplicate genes. Mol Syst Biol. 2010;6:429. Epub 2010/11/18. doi: 10.1038/msb.2010.82. PubMed PMID: 21081923; PubMed Central PMCID: PMCPMC3010121.

82. Gu X, Zhang Z, Huang W. Rapid evolution of expression and regulatory divergences after yeast gene duplication. Proc Natl Acad Sci U S A. 2005;102(3):707–12. Epub 2005/01/14. doi: 10.1073/pnas.0409186102. PubMed PMID: 15647348; PubMed Central PMCID: PMCPMC545572.

83. Kafri R, Bar-Even A, Pilpel Y. Transcription control reprogramming in genetic backup circuits. Nat Genet. 2005;37(3):295–9. Epub 2005/02/22. doi: 10.1038/ng1523. PubMed PMID: 15723064.

84. DeLuna A, Springer M, Kirschner MW, Kishony R. Need-based up-regulation of protein levels in response to deletion of their duplicate genes. PLoS Biol. 2010;8(3):e1000347. Epub 2010/04/03. doi: 10.1371/journal.pbio.1000347. PubMed PMID: 20361019; PubMed Central PMCID: PMCPMC2846854.

85. van der Lee R, Lang B, Kruse K, Gsponer J, Sanchez de Groot N, Huynen MA, et al. Intrinsically disordered segments affect protein half-life in the cell and during evolution. Cell Rep. 2014;8(6):1832–44. Epub 2014/09/16. doi: 10.1016/j.celrep.2014.07.055. PubMed PMID: 25220455; PubMed Central PMCID: PMCPMC4358326.

86. Stelling J, Sauer U, Szallasi Z, Doyle FJ, 3rd, Doyle J. Robustness of cellular functions. Cell. 2004;118(6):675–85. Epub 2004/09/17. doi: 10.1016/j.cell.2004.09.008. PubMed PMID: 15369668.

87. Kitano H. Biological robustness. Nat Rev Genet. 2004;5(11):826–37. Epub 2004/11/03. doi: 10.1038/nrg1471. PubMed PMID: 15520792.

88. Zhou B, Der CJ, Cox AD. The role of wild type RAS isoforms in cancer. Semin Cell Dev Biol. 2016;58:60–9. Epub 2016/07/17. doi: 10.1016/j.semcdb.2016.07.012. PubMed PMID: 27422332; PubMed Central PMCID: PMCPMC5028303.

89. Brachmann CB, Davies A, Cost GJ, Caputo E, Li J, Hieter P, et al. Designer deletion strains derived from Saccharomyces cerevisiae S288C: a useful set of strains and plasmids for PCR-mediated gene disruption and other applications. Yeast. 1998;14(2):115–32.

90. Dobin A, Davis CA, Schlesinger F, Drenkow J, Zaleski C, Jha S, et al. STAR: ultrafast universal RNA-seq aligner. Bioinformatics. 2013;29(1):15–21. Epub 2012/10/30. doi: 10.1093/bioinformatics/bts635. PubMed PMID: 23104886; PubMed Central PMCID: PMCPMC3530905.

91. Patro R, Duggal G, Love MI, Irizarry RA, Kingsford C. Salmon provides fast and bias-aware quantification of transcript expression. Nat Methods. 2017;14(4):417–9. Epub 2017/03/07. doi: 10.1038/nmeth.4197. PubMed PMID: 28263959; PubMed Central PMCID: PMCPMC5600148.

92. Love MI, Huber W, Anders S. Moderated estimation of fold change and dispersion for RNA-seq data with DESeq2. Genome Biol. 2014;15(12):550. Epub 2014/12/18. doi: 10.1186/s13059-014-0550-8. PubMed PMID: 25516281; PubMed Central PMCID: PMCPMC4302049.

93. Love MI, Anders S, Kim V, Huber W. RNA-Seq workflow: gene-level exploratory analysis and differential expression. F1000Res. 2015;4:1070. Epub 2015/12/18. doi: 10.12688/f1000research.7035.1. PubMed PMID: 26674615; PubMed Central PMCID: PMCPMC4670015.

94. Li YY, Douillet C, Huang M, Beck R, Sumner SJ, Styblo M. Exposure to inorganic arsenic and its methylated metabolites alters metabolomics profiles in INS-1 832/13 insulinoma cells and isolated pancreatic islets. Arch Toxicol. 2020;94(6):1955–72. Epub 2020/04/12. doi: 10.1007/s00204-020-02729-y. PubMed PMID: 32277266.

95. Zelena E, Dunn WB, Broadhurst D, Francis-McIntyre S, Carroll KM, Begley P, et al. Development of a robust and repeatable UPLC-MS method for the long-term metabolomic study of human serum. Anal Chem. 2009;81(4):1357–64. Epub 2009/01/28. doi: 10.1021/ac8019366. PubMed PMID: 19170513.

96. Dunn WB, Broadhurst D, Begley P, Zelena E, Francis-McIntyre S, Anderson N, et al. Procedures for large-scale metabolic profiling of serum and plasma using gas chromatography and liquid chromatography coupled to mass spectrometry. Nat Protoc. 2011;6(7):1060–83. Epub 2011/07/02. doi: 10.1038/nprot.2011.335. PubMed PMID: 21720319.

